# Oncogenic Mutant p53 Sensitizes Non-Small Cell Lung Cancer Cells to Proteasome Inhibition via Oxidative Stress-Dependent Induction of Mitochondrial Apoptosis

**DOI:** 10.1101/2024.02.22.581532

**Authors:** Kranthi Kumar Chougoni, Victoria Neely, Boxiao Ding, Eziafa Oduah, Vianna Lam, Bin Hu, Jennifer E. Koblinski, Bradford E. Windle, Swati Palit Deb, Sumitra Deb, Senthil K. Radhakrishnan, Hisashi Harada, Steven R. Grossman

**Affiliations:** Department of Medicine, Keck School of Medicine of USC and USC Norris Comprehensive Cancer Center, University of Southern California, Los Angeles, CA USA; Philips Institute for Oral Health Research, VCU Schools of Dentistry and Medicine, Virginia Commonwealth University, Richmond, VA USA; Department of Internal Medicine, VCU Schools of Dentistry and Medicine, Virginia Commonwealth University, Richmond, VA USA; VCU Cancer Mouse Models Core, VCU Schools of Dentistry and Medicine, Virginia Commonwealth University, Richmond, VA USA; VCU Massey Comprehensive Cancer Center, Departments of VCU Schools of Dentistry and Medicine, Virginia Commonwealth University, Richmond, VA USA; Pathology, and VCU Schools of Dentistry and Medicine, Virginia Commonwealth University, Richmond, VA USA; Biochemistry & Molecular Biology, VCU Schools of Dentistry and Medicine, Virginia Commonwealth University, Richmond, VA USA

## Abstract

Non-small cell lung cancer (NSCLC) cells with oncogenic mutant p53 alleles (Onc-p53) exhibit significantly higher levels of proteasome activity, indicating that Onc-p53 induces proteotoxic stress which may be leveraged as a therapeutic vulnerability. Proteasome inhibitors (PIs), such as bortezomib (BTZ), can induce toxic levels of oxidative stress in cancer cells and thus we investigated whether PIs exhibit preferential cytotoxicity in Onc-p53 NSCLC cells. Indeed, BTZ and other PIs exhibited the IC_50_ 6-7-fold lower in Onc-p53 cells vs. wild-type (WT) p53 cells. BTZ cytotoxic effects in Onc-p53 cells were nearly completely rescued by antioxidants such as N-acetyl cysteine, indicating that oxidative stress is the critical driver of BTZ-dependent cytotoxic effects in Onc-p53 cells. Importantly, we observed oxidative stress-dependent transcriptional induction of the pro-apoptotic NOXA with downstream cleaved caspase-3, consistent with apoptotic cell death in Onc-p53 but not in WT p53 cells treated with BTZ, and BTZ-generated oxidative stress was linked to nuclear translocation of NRF2 and transcriptional activation of ATF3, which in turn was required for NOXA induction. Validating BTZ’s translational potential in Onc-p53 NSCLC, BTZ and carboplatin or the BH3-mimetic navitoclax were synergistically cytotoxic in Onc-p53 but not WT p53 cells *in vitro,* and BTZ effectively limited growth of Onc-p53 NSCLC xenografts when combined with either carboplatin or navitoclax *in vivo*. Our data therefore support further investigation of the therapeutic utility of PIs combined with carboplatin or BH3-mimetics in Onc-p53 human NSCLC as novel therapeutic strategies.

**Significance:** Non-small cell lung cancer (NSCLC) is the leading cause of cancer death due, in part, to a lack of active therapies in advanced disease. We demonstrate that proteasome inhibitor/BH3-mimetic combination therapy is an active precision therapy in NSCLC cells and tumors expressing oncogenic mutant p53 alleles (Onc-p53).

## Introduction

Lung Cancer is one of the most commonly diagnosed cancer in men and women and non-small cell lung cancer (NSCLC) is the most common histologic subtype of human lung cancer (84%), which is the leading cause of cancer-related death annually in the US (1,2). The molecular underpinnings of lung oncogenesis are becoming better understood, and recent evidence indicates that missense mutation of the p53 tumor suppressor, which occurs in nearly 70% of NSCLC tumors, creates emergent oncogenic activities of the Onc-p53 alleles that drive enhanced proliferation, metastasis and treatment resistance, consistent with worse outcomes observed in patients with Onc-p53 NSCLC (3–6). However, as Onc-p53 loses all functions associated with native WT p53, Onc-p53 alleles offer the potential to exploit associated cancer cell-specific vulnerabilities for therapeutic benefit (7,8).

One such potential vulnerability is the overexpression of 26S proteasome subunits and excess proteasome activity consistent with proteotoxic stress that has been observed in many tumor cell types expressing Onc-p53, including lung cancer (9,10). Proteasome inhibitors (PIs) induce cellular stress at multiple levels, including proteotoxic, autophagic and oxidative stress, but are particularly effective in neoplastic cells with ongoing proteotoxic stress (11). However, the clinical utility of the PI drug class has been limited by lack of efficacy outside of B cell malignancies, such as multiple myeloma (MM) and mantle cell lymphoma, which are uniquely sensitive to PIs due to immunoglobulin-synthesis related proteotoxic stress (12).

The mechanism of PI resistance of most solid tumors, including NSCLC, remains unclear, as there has been limited exploration of precision molecular techniques that might identify subsets of tumors sensitive to PI alone or combined with other targeted agents, as has been successfully deployed in MM. Relevant to known mechanisms of PI cytotoxic action, an additional vulnerability associated with Onc-p53 expression in many solid tumor types is the presence of excess reactive oxygen species (ROS) (13). Excess ROS in the baseline state for cancer cells could theoretically sensitize them to therapeutics such as PIs that generate additional ROS to a level toxic to cancer but not normal cells due to additional deficiencies in ROS scavenging mechanisms (14,15).

Based on previously described overexpression of proteasome subunits and excess proteasome activity in NSCLC cells expressing Onc-p53 (9), we investigated whether PIs exhibit preferential cytotoxicity in Onc-p53 cells *in vitro* and *in vivo*. Indeed, bortezomib (BTZ) and other PIs were preferentially cytotoxic in Onc-p53 expressing NSCLC cells which also exhibited higher levels of proteasome activity, indicating that Onc-p53 induces proteotoxic stress. Importantly, we observed an oxidative stress-dependent transcriptional cascade in cells treated with BTZ, resulting in induction of the pro-apoptotic BH3-only protein NOXA and apoptotic cell death. Validating BTZ’s translational potential in Onc-p53 NSCLC, combinations of BTZ with carboplatin or the BH3-mimetic navitoclax were synergistically cytotoxic in Onc-p53 *in vitro,* and effectively limited growth of Onc-p53-expressing NSCLC xenografts *in vivo*. Our data therefore support further investigation of the therapeutic utility of PIs combined with carboplatin or BH3-mimetics in Onc-p53 human NSCLC as novel therapeutic strategies for many NSCLC patients who do not benefit from targeted or immunotherapies.

## Results

### NSCLC cells expressing Onc-p53 exhibit enhanced proteasome activity and sensitivity to proteasome inhibitors

Based on findings in multiple other solid tumors that Onc-p53 elevates proteasome subunit gene expression and activity (9), we hypothesized that NSCLC cells expressing Onc-p53 might also demonstrate similar upregulation of proteasome activity, and as a result, also exhibit enhanced cytotoxic sensitivity toward PIs. Therefore, we first assayed 20S proteasome activity in a panel of NSCLC cell lines with differing p53 mutational status (**Fig. 1A**). As seen in other solid tumor cell types (9), Onc-p53 expressing cells demonstrated 3∼4-fold higher 20S proteasome activity than WT p53 expressing cells (*p*<0.05; **Fig. 1A**). To determine if Onc-p53 was necessary for the upregulation of proteasome activity, we developed H1975-Control and homozygous p53 knockout (H1975-p53KO) cells using CRISPR/Cas9 technology (**Fig. 1B, right**). As has been observed in breast cancer cells (9), 20S proteasome activity was strongly suppressed when Onc-p53 was deleted (**Fig. 1B, left**). These results indicate that Onc-p53 is required for the increased 20S proteasome activity found in Onc-p53 NSCLC cells.

**Fig. 1.**
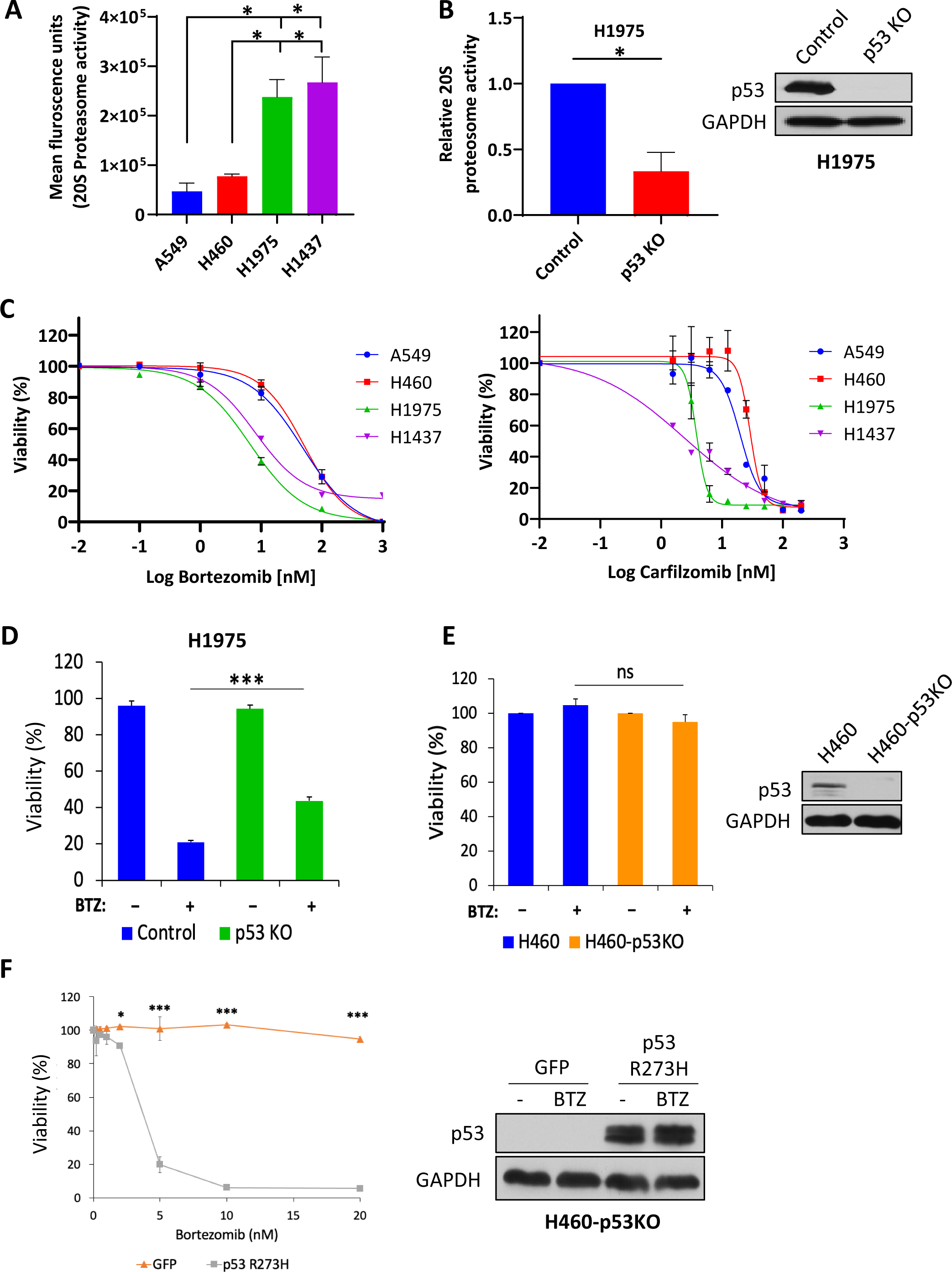
Onc-p53 regulates basal levels of 20S proteasome activity and is required for BTZ-induced cytotoxicity in NSCLC cells. **A.** 20S proteasome activity was determined in the indicated cell lines using a fluorometric proteosome activity assay. All pairwise comparisons were made using student’s t-test. **B. (Left)** Basal levels of 20S proteasome activity in lysates of H1975 Control and p53 KO cells. (**Right**) Cell lysates of H1975-Control and H1975-p53KO (p53KO) cell lines generated using CRISPR/Cas9 were immunoblotted with p53 and GAPDH antibodies. **C.** The indicated cells were treated with vehicle or increasing concentrations of bortezomib (BTZ; left) or carfilzomib (CFZ; right) for 72 h, and cell viability was determined by the MTT assay. **D**. H1975-Control and H1975-p53KO cells were treated with vehicle (-) or BTZ (5 nM) for 48 h and cell viability was determined by Trypan-blue exclusion assay. **E.** (**Left**) H460 and H460-p53KO cells were treated with vehicle (-) or BTZ (5 nM) for 72 h and cell viability was determined by WST-1 assay. (**Right**) Cell lysates of H460 and H460-p53KO cells (16) were immunoblotted with p53 and GAPDH antibodies. **F.** (**Left**) H460-p53KO cells stably expressing GFP or p53R273H cDNA were treated with increasing concentrations of BTZ for 72 h and cell viability was determined by WST-1 assay. (**Right**) Cell lysates of H460-p53KO cells stably expressing GFP or p53R273H cDNA were immunoblotted with p53 and GAPDH antibodies. \**p*<0.05, \*\**p*<0.01, \*\*\**p*<0.005, ns indicates *p*>0.05. Error bars indicate +/- 1.0 S.D.

As the increased 20S proteasome activity in Onc-p53 NSCLC cells might signal enhanced sensitivity to PIs, we explored PI cytotoxicity in Onc-p53 vs. WT p53 NSCLC cells. Indeed, H1975 (p53^R273H^) and H1437 (p53^R267P^) Onc-p53 cells were much more sensitive to proteasome inhibition by BTZ (IC_50_ 7 nM and 8 nM, respectively) than NSCLC cell lines harboring WT p53, including A549 (IC_50_ 49 nM) and H460 (IC_50_ 51 nM) (**Fig. 1C, left; Table 1**). This differential sensitivity to proteasome inhibition between Onc-p53 and WT p53 cells was also manifested using another FDA-approved PI, carfilzomib (CFZ) (**Fig. 1C, right; Table 1**).

To determine if Onc-p53 was necessary for the enhanced sensitivity of Onc-p53 cells to BTZ-induced cell death, we tested H1975-Control and H1975-p53KO cells for BTZ sensitivity and observed that H1975-p53KO cells were significantly less sensitive to BTZ treatment as compared with H1975-Control cells (60% vs. 80% loss of viability, respectively; *p*<0.01; **Fig. 1D**). In contrast, H460 cells with CRISPR knockout of the WT p53 gene (H460-p53KO; **Fig. 1E, right panel**) (16) did not show sensitization to BTZ treatment relative to parental H460 cells (<5% loss of viability in both cell lines; **Fig. 1E, left panel**). However, expression of exogenous p53^R273H^ cDNA vs. GFP control in H460 KO cells (**Fig. 1F, right panel**) robustly restored BTZ cytotoxicity (IC_50_ 4 nM) (**Fig. 1F, left panel**). These findings suggest that Onc-p53 induces proteasome activity and renders NSCLC cells differentially more vulnerable to PI treatment.

### Onc-p53 attenuates glutathione levels in NSCLC cells

Considering that the mechanism of BTZ cytotoxicity often involves induction of reactive oxygen species in sensitive cancer cells (15) and that Onc-p53 might alter tumor cell oxidative state (13), we mined the metabolomic profiles of 52 human NSCLC cell lines that included those expressing Onc-p53 alleles vs. those exhibiting p53 loss-of-function (LOF; homozygous deletion, frameshift and early termination alleles associated with loss of heterozygosity) for abundance of the key antioxidant metabolite glutathione (both reduced and oxidized species of glutathione – GSH and GSSG, respectively) (17). The average GSH/GSSG levels were ∼50% lower in Onc-p53 NSCLC cell lines (*n* = 41) compared to the LOF p53 cell lines (*n* = 11) (*p*<0.01; **Fig. 2A**). We then measured total GSH levels in the WT and Onc-p53 NSCLC cell lines from **Fig. 1A** and observed that the average levels of GSH in Onc-p53 expressing cell lines were significantly lower than in WT p53 cell lines (*p*<0.05; **Fig. 2B**). Furthermore, loss of Onc-p53 in H1975-p53KO vs. H1975-Control cells led to a 70% increase in GSH levels (*p*<0.01; **Fig. 2C**). Thus, Onc-p53 expression is associated with glutathione depletion in NSCLC cells.

**Fig. 2.**
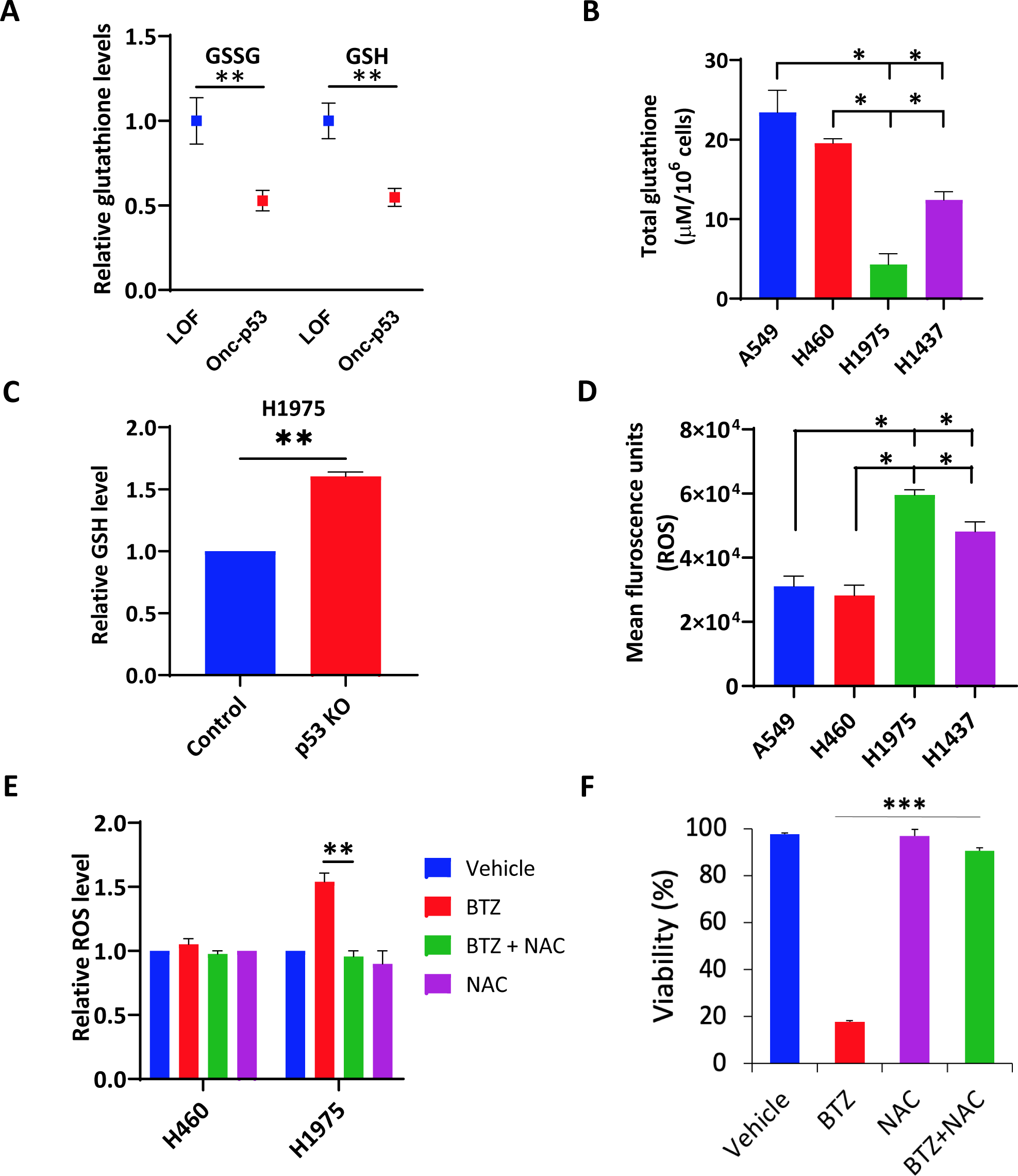
Onc-p53 NSCLC cells exhibit basal and PI-induced oxidative stress. **A.** Total glutathione (GSSG & GSH) profiling of NSCLC cell lines that either express oncogenic mutant p53 alleles (Onc-p53) or lack p53 protein altogether due to either homozygous deletion or frameshift/premature termination mutations (LOF). Data were analyzed using unpaired *t* test. **B-C.** Total (**B**) or relative (**C;** H1975-p53KO glutathione level normalized to glutathione level in H1975-Control) glutathione levels were determined in the indicated cell lines. **D.** Basal ROS levels were estimated in the indicated cell lines using a fluorescent ROS detection assay and plotted as Mean Fluorescence Units. **E.** H460 or H1975 cells were treated with vehicle, BTZ (5 nM), N-acetylcysteine (NAC; 1 mM) or BTZ + NAC for 24 h. ROS was measured by staining with DCFH-DA using flow cytometry. **F.** H1975 cells were treated as per **E** for 48 h and cell viability was assessed by Trypan-blue excursion. All pairwise comparisons were made using student’s t-test. \**p*< 0.05, \*\**p*< 0.01, \*\*\**p*<0.005. Error bars indicate +/- 1.0 S.D.

### Induction of ROS by BTZ requires Onc-p53

Based on the depletion of glutathione levels seen with Onc-p53 expression, we inferred that the presence of Onc-p53 might lead to higher levels of basal ROS as well as potentially toxic accumulation of ROS upon exposure to BTZ. Indeed, H1975 and H1437 Onc-p53 cells demonstrated significantly higher average levels of basal ROS than H460 and A549 WT p53 cells as assessed by fluorescent ROS indicator dye (∼2-fold increase, *p*<0.05; **Fig. 2D**). We then explored whether intracellular ROS level contributes to enhanced BTZ cytotoxicity in WT vs. Onc-p53 cells by staining BTZ-, or BTZ + ROS scavenger *N*-acetyl-L-cysteine (NAC)-treated H1975 vs. H460 cells with ROS indicator dye followed by flow cytometry. Interestingly, BTZ significantly increased ROS levels in Onc-p53 H1975 cells (*p*<0.01) but not in WT p53 H460 cells, and inclusion of NAC reversed the increase in ROS in H1975 cells (**Fig. 2E**). Furthermore, NAC as well as the GSH-mimetic, glutathione ethyl ester (GSH-EE) nearly completely rescued BTZ-induced cell death in H1975 cells (*p*<0.005 and *p*<0.05, respectively; **Figs. 2F and S1A**), suggesting that the levels of ROS scavengers, such as glutathione, were relatively deficient, allowing accumulation of toxic levels of ROS after BTZ treatment. These results collectively suggest that the BTZ sensitivity of Onc-p53 NSCLC cells is due to the induction of a toxic level of ROS not seen in NSCLC cells expressing WT p53.

### BTZ induces NOXA and caspase-dependent apoptosis

To determine the mode of action of BTZ-induced cell death in Onc-p53-expressing NSCLC cells, we first determined the contribution of apoptosis by testing the effect of the pan-caspase inhibitor, QVD-OPh in BTZ-induced cell death of H1975 cells, and remarkably, cell death was almost completely rescued by QVD-OPh (*p*<0.005; **Fig. 3A**). BTZ also induced cell surface exposure of the apoptotic marker Annexin V in nearly 80% of cells at 48 h as measured by flow cytometry, which was partially rescued by QVD-OPh (*p*<0.005; **Fig. 3B**). Moreover, the apoptosis marker cleaved caspase-3 was induced by BTZ in H1975 cells, with complete rescue of caspase-3 cleavage by QVD-OPh (**Fig. 3C**). Taken together, these data strongly suggest that BTZ-induced cell death is largely mediated by apoptosis.

**Fig. 3.**
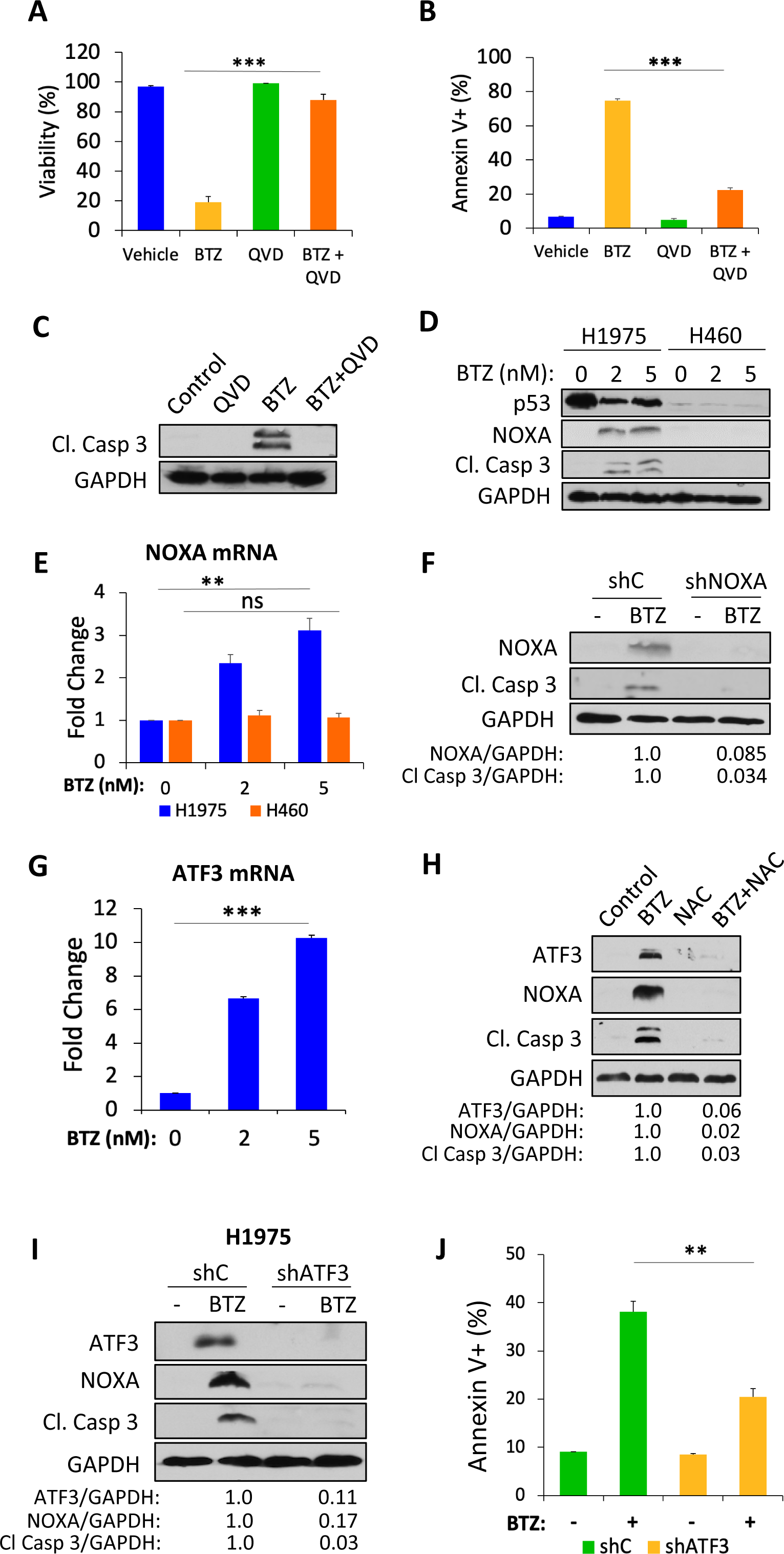
BTZ induces ATF3/NOXA and caspase-dependent apoptosis. **A.** H1975 cells were pre-treated (30 min) with vehicle or the pan-caspase inhibitor QVD-OPh (QVD; 1 µM), then treated with vehicle or BTZ (5 nM) for 48 h. Cell viability was determined by Trypan-blue exclusion assay. **B.** H1975 cells were treated as in **A** and percentage of apoptotic cells was determined by Annexin V-PI staining followed by FACS analysis. **C.** Cell lysates from **A** were immunoblotted with the indicated antibodies. **D.** H1975 or H460 cells were treated with vehicle, 2, or 5 nM of BTZ for 48 h and cell lysates were immunoblotted with the indicated antibodies. **E.** RNA from **D** was extracted and subjected to qRT-PCR with NOXA primers. **F.** H1975 cells stably expressing control shRNA (shC) or NOXA shRNA (shNOXA) were treated with vehicle or BTZ (5 nM) for 48 h and cell lysates were immunoblotted with the indicated antibodies. **G.** H1975 cells were treated with vehicle, 2, or 5 nM of BTZ for 48 h and mRNA expression of ATF3 was analyzed by qRT-PCR. **H.** H1975 cells were treated with vehicle, NAC (1 mM), BTZ (5 nM) or BTZ + NAC for 48 h and cell lysates were immunoblotted with indicated antibodies. **I.** H1975 cells stably expressing control shRNA (shC) or ATF3 shRNA (shATF3) were treated with vehicle or BTZ (5 nM) for 48 h and cell lysates were immunoblotted with the indicated antibodies. **J.** H1975 cells stably expressing control shRNA (shC) or ATF3 shRNA (shATF3) were treated with vehicle or BTZ (5 nM) for 48 h and the percentage of apoptotic cells was determined by Annexin V-PI staining followed by FACS analysis. \*\**p*< 0.01, \*\*\**p*<0.005, ns indicates *p*>0.05. Error bars indicate +/- 1.0 S.D.

Given that the mechanism of BTZ-induced cell death of Onc-p53 H1975 cells is apoptotic, as is also seen in MM cells (18), we therefore investigated whether the expression of the BH3-only pro-apoptotic protein NOXA, which is critical for BTZ-induced apoptosis in MM cells (18), was also induced in NSCLC cells. Indeed, NOXA protein expression was strongly induced after BTZ exposure in both H1975 and H1437 Onc-p53 cells, but not H460 WT p53 cells (**Figs. 3D and S1B**). Furthermore, the induction of NOXA protein in H1975, but not H460, cells was mirrored by *NOXA* mRNA expression, suggesting BTZ regulates NOXA primarily via transcription (**Fig. 3E**).

To explore the requirement of NOXA for BTZ-induced apoptosis in H1975 cells, we stably expressed shNOXA or scrambled shRNA in H1975 and H1437 cells and exposed both cell lines to vehicle or BTZ (**Figs. 3F, S1C-D**). Whereas NOXA and cleaved caspase-3 were induced by BTZ in cells expressing control shRNA, silencing of NOXA resulted in significant reduction of cleaved caspase-3 levels after BTZ treatment (**Figs. 3F, S1C-D**), reflective of a requirement for NOXA for BTZ-induced apoptosis in Onc-p53 expressing H1975 and H1437 cells.

### ATF3 mediates oxidative stress-dependent induction of NOXA by BTZ in Onc-p53 NSCLC cells

As NOXA induction by BTZ occurred at the level of mRNA expression, we next investigated possible factor(s) controlling NOXA transcription in the setting of BTZ exposure. Though NOXA can be transcriptionally regulated by numerous transcription factors, ATF3 and ATF4 are both known to regulate NOXA in the setting of cellular stress (19), and we noted that transcription of *ATF3* mRNA was strongly induced in BTZ vs. vehicle-treated H1975 cells (**Fig. 3G**), while *ATF4* mRNA levels did not change (**Fig. S2A**). Analysis of ATF3 protein expression likewise revealed strong induction, along with NOXA and cleaved caspase-3 upon BTZ treatment of H1975 or H1437 cells (**Figs. 3H, S1B, and S2B**). Consistent with a critical role for oxidative stress in BTZ induction of cell death (**Figs. 2F and S1A**), both NAC and GSH-EE abrogated the induction of ATF3, NOXA and cleaved caspase-3 by BTZ in H1975 cells (**Figs. 3H**, **S2C-G**), indicating that oxidative stress is a critical factor downstream of BTZ in activating the signaling cascade in Onc-p53 cells that ultimately leads to cell death. We next determined whether ATF3 is required for BTZ induction of NOXA and apoptosis in Onc-p53 NSCLC cells, by interrogating NOXA and cleaved caspase-3 induction in BTZ-treated H1975 cells expressing ATF3 vs. control shRNA. Indeed, relative to H1975 cells stably expressing control shRNA, NOXA and cleaved caspase-3 induction was attenuated upon BTZ treatment in H1975 clones stably expressing two independent shATF3 constructs (**Figs. 3I and S2H)**. Moreover, ATF3 knockdown led to a 50% reduction in the Annexin V-positive apoptotic cell population upon BTZ exposure (*p*<0.01; **Fig. 3J**). These results suggest that BTZ initiates an oxidative stress-dependent signaling cascade in Onc-p53 NSCLC cells where ATF3 transcriptional induction downstream of BTZ exposure induces NOXA expression, and ultimately, apoptosis.

### BTZ induces nuclear relocalization and accumulation of NRF2 in Onc-p53 expressing NSCLC cells

We next addressed the mechanism by which oxidative stress induced by BTZ might be connected to ATF3/NOXA induction. Of potential regulators of ATF3, NRF2 is a well known transcription factor regulating the cellular antioxidant response, and *ATF3* transcription can be directly regulated by NRF2 (20–22). Under basal unstressed conditions, NRF2 is held in an inactive cytpoplasmic complex with the E3 ubiquitin ligase KEAP1, but upon induction of oxidative cellular stress, NRF2 is released from KEAP1, translocates to the nucleus, and regulates numerous cellular stress-response transcriptional programs (23,24). We thus examined the subcellular localization of NRF2 in vehicle vs. BTZ-treated H1975 cells by immunofluorescence, and observed that NRF2 relocalized from cytosolic/nuclear dual localization to exclusively nuclear localization upon BTZ treatment which was abrogated by NAC, indicating that NRF2 relocalization was likely due to oxidative stress (**Fig. 4A-B**) (24). Furthermore, total cellular NRF2 protein abundance was induced in BTZ vs. vehicle-treated H1975 cells, consistent with reported NRF2 stabilization that occurs upon dissociation from KEAP1 (**Fig. 4C**) (24). NAC treatment reduced the induction of NRF2 abundance consistent with an oxidative stress dependent mechanism regulating NRF2 abundance after BTZ (**Fig. 4C**). Taken together, the abrogation of BTZ-mediated nuclear relocalization and induction of NRF2 abundance by NAC aligns with the inhibition of BTZ-mediated ATF3/NOXA/cleaved caspase-3 induction and loss of cell viability by NAC and GSH-EE (**Figs. 2F, 3H, S1A, S2C-G**), suggesting that NRF2 could be the factor connecting BTZ-induced oxidative stress to induction of ATF3/NOXA and apoptosis in Onc-p53 NSCLC cells.

**Fig. 4.**
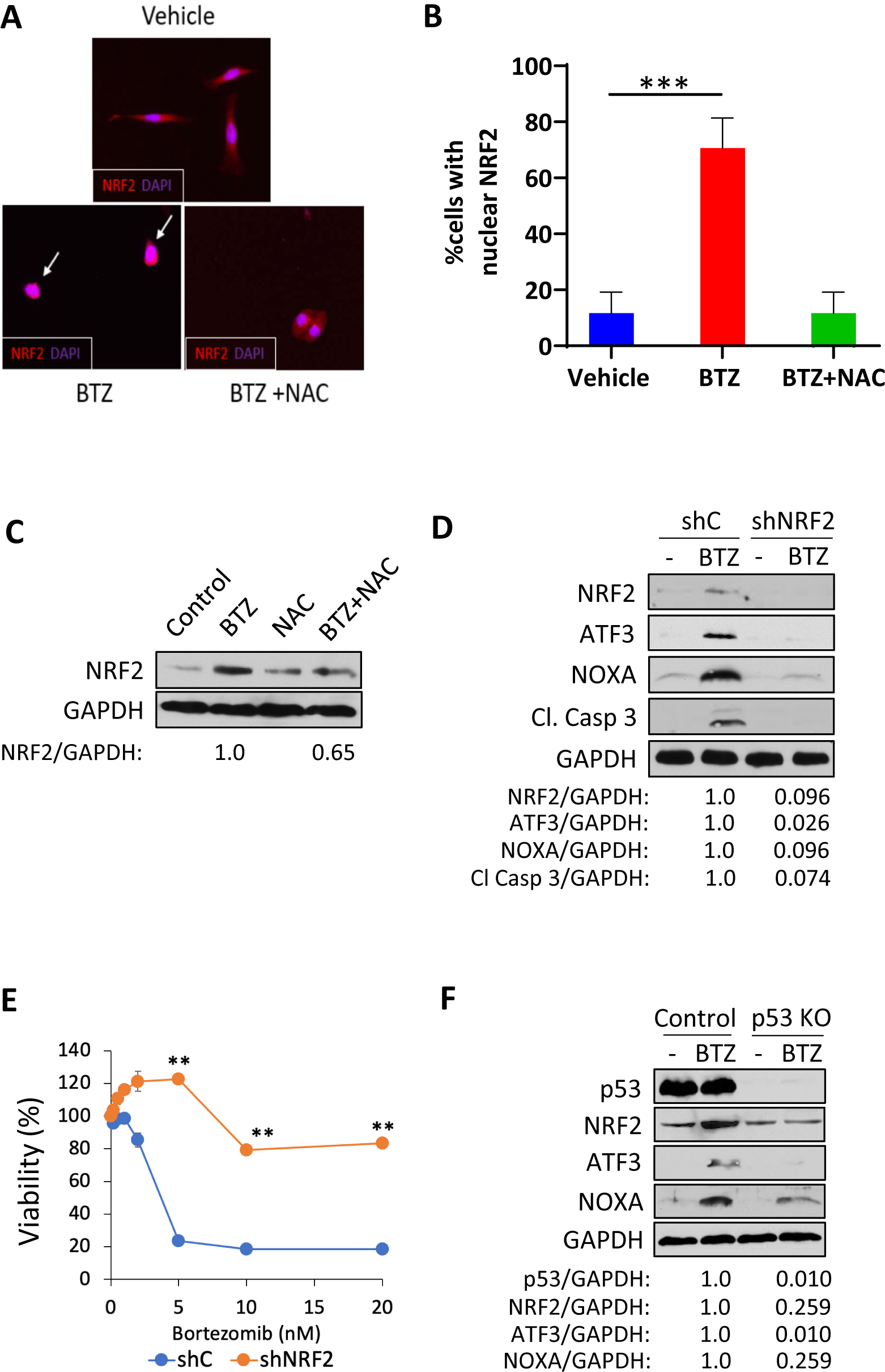
The Onc-p53-NRF2-ATF3-NOXA pathway contributes to BTZ-induced apoptosis in Onc-p53 NSCLC cells. **A.** H1975 cells were treated with vehicle or BTZ (10 nM) ± NAC (1 mM) for 24 h and immunofluorescence was performed with NRF2 antibody (red) and DAPI (blue). Merged images of NRF2 and DAPI stains are shown. **B.** Quantification of the percentage of cells with NRF2 localization exclusive to the nucleus from 10 random fields (>10 cells/field counted) at 20X magnification. A pairwise comparison was made using student’s t-test between the two groups. **C.** H1975 cells were treated with vehicle, NAC (1 mM), BTZ (5 nM) or BTZ + NAC for 48 h and cell lysates were immunoblotted with NRF2 and GAPDH antibodies. **D.** H1975 cells stably expressing control shRNA (shC) or NRF2 shRNA (shNRF2) were treated with vehicle or BTZ (5 nM) for 48 h and cell lysates were immunoblotted with indicated antibodies. **E.** H1975 shRNA control and shNRF2 cells were treated with vehicle or the indicated concentrations of BTZ for 72 h and cell viability was determined by WST-1 assay. **F.** H1975-Control or H1975-p53 KO cells were treated with vehicle (-) or BTZ (5 nM) for 48 h, followed by immunoblotting of cell lysates with indicated antibodies. \*\**p*<0.01, \*\*\**p*<0.005. Error bars indicate +/- 1.0 S.D.

### NRF2 is required for induction of ATF3/NOXA and cell death by BTZ in Onc-p53 expressing NSCLC cells

To determine whether NRF2 is required upstream of BTZ-mediated cytotoxic signaling via ATF3/NOXA in Onc-p53 NSCLC cells, we first treated H1975 cells stably expressing NRF2 or control shRNA with vehicle or BTZ, and immunoblotted cell lysates for NRF2, ATF3, NOXA, and cleaved caspase-3 (as a surrogate for apoptosis; **Fig. 4D**). While, BTZ treatment of shControl-expressing H1975 cells revealed the expected induction of ATF3/NOXA and cleaved caspase-3, shNRF2 expression strongly attenuated BTZ induction of ATF3/NOXA and caspase-3 cleavage (**Fig. 4D**). Moreover, shNRF2 expression rendered H1975 cells highly resistant to BTZ-mediated cell death, as only 20% loss of viability was observed at BTZ doses as high as 20 nM, while shControl-expressing H1975 cells exhibited expected sensitivity to BTZ with ∼80% loss of viability with exposure to only 5 nM BTZ (*p*<0.01; **Fig. 4E**). These results indicate that NRF2 is necessary for BTZ induction of ATF3/NOXA and apoptosis in Onc-p53 H1975 NSCLC cells.

### Onc-p53 is required for BTZ-mediated induction of NRF2/ATF3/NOXA and apoptosis in Onc-p53 expressing NSCLC cells

We next addressed if activation of the proposed NRF2-ATF3-NOXA signaling cascade in BTZ-treated cells ultimately requires ongoing expression of Onc-p53. Indeed, H1975-p53KO cells, which exhibit resistance to BTZ-mediated cytotoxic cell death (**Fig. 1D**), demonstrated reduced or nearly absent induction of NRF2, ATF3 and NOXA protein as well as *ATF3* and *NOXA* mRNA (**Figs. 4F, S3A-B**). These results indicate that Onc-p53 is a necessary upstream component of an NRF2-ATF3-NOXA signaling cascade that leads to apoptosis in Onc-p53 H1975 cells.

### BTZ/carboplatin combination limits tumor growth of Onc-p53-harboring NSCLC xenografts *in vivo*

The standard of care chemotherapeutic agent, carboplatin is used routinely in both early stage and advanced NSCLC. We thus tested the efficacy of BTZ in combination with carboplatin in NSCLC cells. We found that BTZ enhanced carboplatin-mediated cytotoxicity in a dose-dependent fashion in Onc-p53 H1975 cells (*p*<0.01; **Fig. 5A**), but not in WT p53 A549 cells (**Fig. 5B**) *in vitro*. To verify these findings *in vivo*, we treated immunocompromised mice bearing H1975 or A549 subcutaneous xenografts for 4 weeks with vehicle, BTZ, carboplatin, or the combination (**Fig. 5C**). In H1975 xenografted mice, BTZ or carboplatin by themselves showed modest but not statistically significant reductions in tumor volume compared with vehicle treatment, whereas the BTZ/carboplatin combination significantly reduced tumor volume by ∼50% as compared to vehicle treatment (*p*<0.001; **Fig. 5C**). However, none of the treatments reduced tumor volume at the experimental endpoint in A549 xenografted mice (**Fig. 5D**). These results suggest that the *in vivo* impact of BTZ on therapy of Onc-p53 NSCLC tumors (with R273H allele as proof of principle) can be improved in a cooperative manner by a platinum-chemotherapy agent that is already in use as a standard of care in NSCLC and is also known to induce oxidative stress (25).

**Fig. 5.**
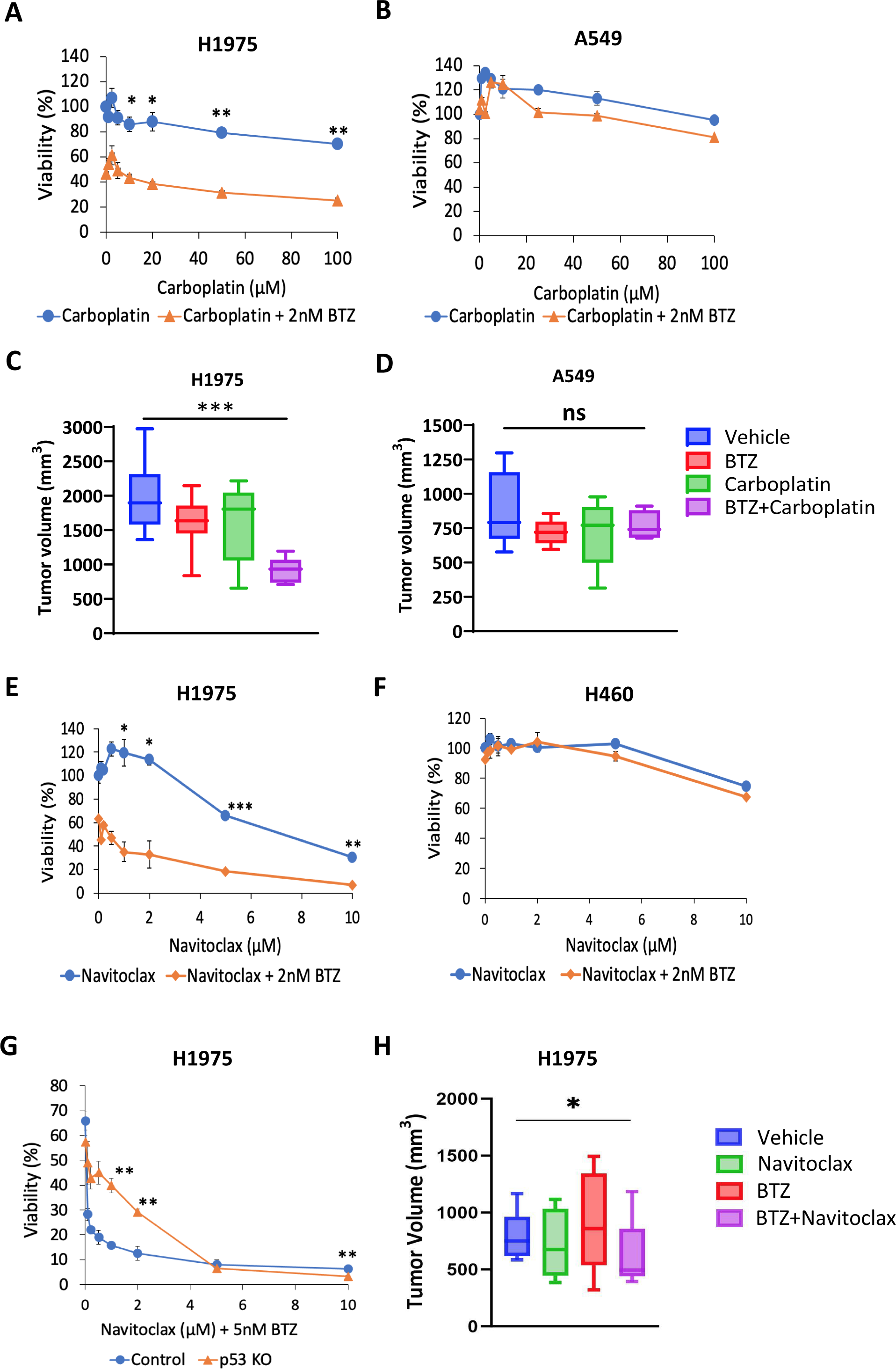
Carboplatin and navitoclax enhance BTZ-induced cytotoxicity. **A-B.** H1975 and H460 cells were treated with vehicle or the indicated concentrations of carboplatin with/without BTZ (2 nM) for 96 h. Cell viability was determined by WST-1 assay. **C-D.** NSG mice bearing H1975 or A549 xenografts were treated with vehicle, BTZ, carboplatin (CBP), or BTZ/carboplatin combination for 4 weeks. Tumor volume was measured at the end of the study (N = 10). Error bars indicate SEM. **E-F.** H1975 and H460 cells were treated with vehicle or the indicated concentrations of navitoclax with/without BTZ (2 nM) for 96 h. Cell viability was determined by WST-1 assay. **G.** H1975-Control or H1975-p53KO cells were treated with vehicle or the indicated concentrations of navitoclax with/without BTZ (5 nM) for 72 h. Cell viability was determined by WST-1 assay. **H.** NSG mice were injected with H1975 cells (4.0 x 10^6^) and once tumors became palpable, mice were treated for 4 weeks with vehicle (control), navitoclax (80 mg/kg) 3x/week via oral gavage, BTZ (0.6 mg/kg) 2x/week via intraperitoneal injection, or the BTZ + navitoclax combination. Tumor volume was measured at the end of the study (N = 4/group). Error bars indicate SEM. \**p*<0.05, \*\**p*<0.01, \*\*\**p*<0.005, ns indicates *p*>0.05. Error bars indicate +/- 1.0 S.D., except where indicated as SEM.

### BH3 mimetics sensitize Onc-p53-harboring NSCLC cells to BTZ cytotoxicity

To further explore additional mechanisms of specifically enhancing PI cytotoxicity in Onc-p53 NSCLC cells, we considered whether targeting BCL-2-family anti-apoptotic proteins might cooperatively improve BTZ cytotoxicity as well. First, we investigated the basal levels of anti-apoptotic BCL-2 family proteins in H1975 and H1437 Onc-p53 cells and observed that BCL-X_L_ and MCL-1 were variably expressed in either cell line, but BCL-2 was undetectable in H1437 cells (**Fig. S4A**). As NOXA specifically binds to and inactivates pro-survival MCL-1 (26), we hypothesized that the BH3-mimetics targeting BCL-X_L_, such as navitoclax (ABT-263, BCL-2/BCL-X_L_ dual-inhibitor) could enhance BTZ-induced cytotoxicity. Indeed, we found that BTZ strikingly enhanced the cytotoxicity of navitoclax in Onc-p53 H1975 and H1437 cells (*p*<0.05; **Figs. 5E and S4B**), but not in WT p53 H460 cells (**Fig. 5F**), and H1975-p53KO cells were much less sensitive to the combination treatment as compared with H1975-Control cells (*p*<0.01; **Fig. 5G**), indicating Onc-p53 dependency.

These striking *in vitro* data suggest that the combination of BTZ and a BH3-mimetic could represent an effective rational combination therapeutic strategy in NSCLC bearing Onc-p53. As such, we next tested the efficacy of the combination of BTZ and navitoclax *in vivo* by treating H1975-xenografted NSG mice with vehicle, BTZ, navitoclax, or the combination, over 4 weeks. Relative to the BTZ/carboplatin combination which caused a 50% reduction in final tumor volume relative to vehicle treated tumors, the BTZ/navitoclax combination caused a more modest but still significant ∼30% reduction in tumor volume after 4 weeks of treatment (*p*<0.05; **Fig. 5H**). Consistent with the proposed apoptotic mechanism of BTZ cytotoxicity in NSCLC cells, immunohistochemical staining of the treated tumors for cleaved caspase-3 revealed strongly enhanced staining (>50% of nuclei positive) in tumors exposed to BTZ/navitoclax combination treatment vs. vehicle or either drug alone (**Fig. S4C and S4D**). Notably, the BTZ/navitoclax combination was well-tolerated, despite the known dose-limiting toxicities of navitoclax, including thrombocytopenia, that prevented its further clinical development (27,28). Indeed, platelet and neutrophil counts obtained at the endpoint of the BTZ/navitoclax experiment in **Fig. 5H** actually increased in all treatment groups relative to vehicle-treated mice and remained within the normal range (**Fig. S4E**) (29). These data support further study of combination therapy with PIs and BCL-X_L_ inhibitors for Onc-p53 expressing NSCLC patient tumors.

## Discussion

In the current study, we have identified a vulnerability of NSCLC cells to PIs dependent on Onc-p53 expression which was known to drive excess levels of proteasome gene expression and activity in multiple tumor settings (9). Indeed, we confirmed that Onc-p53 NSCLC cells also exhibit significantly higher levels of proteasome activity requiring the ongoing expression of Onc-p53, and that PIs exhibit preferential cytotoxicity in Onc-p53 vs. WT or cells lacking p53 *in vitro* and *in vivo*. Surprisingly, BTZ cytotoxic effects in Onc-p53 NSCLC cells were rescued completely by NAC, indicating that oxidative stress is a critical driver of BTZ-dependent cytotoxic effects in Onc-p53 cells. Importantly, we observed oxidative stress-dependent nuclear translocation of NRF2 and transcriptional activation of ATF3, which in turn was required for NOXA induction and apoptosis in Onc-p53 NSCLC cells treated with BTZ. Validating BTZ’s translational potential in Onc-p53 NSCLC, BTZ and the standard chemotherapeutic carboplatin or the BH3-mimetic navitoclax were cooperatively, if not synergistically, cytotoxic in Onc-p53 but not WT p53 NSCLC cells *in vitro,* and BTZ effectively limited growth of Onc-p53-expressing NSCLC xenografts when combined with either carboplatin or navitoclax *in vivo*.

Although PIs are clinically approved as part of highly active and life prolonging therapy for the treatment of multiple myeloma, their therapeutic utility in NSCLC, or any other solid tumors, has never been established (30). BTZ and CFZ were previously tested as single agents, or in combination with platinum or other chemotherapeutic agents, in over 20 Phase I/II NSCLC clinical trials (10,31) with a modest efficacy signal observed in a few of the combination trials (with docetaxel, gemcitabine, or carboplatin), mostly in relapsed/refractory disease (32,33). However, gaps in patient selection criteria and other inconsistencies preclude the ability to draw any conclusions on efficacy. For example, a study in which BTZ was used in combination with docetaxel had no significant anti-tumor effect, while some other studies were prematurely terminated due to adverse outcomes in patients, such as thrombocytopenia or neutropenia (33–35). The criteria for the patients included in these trials varied from one trial to the other and was not biomarker dependent (10). Our *in vitro/in vivo* results strongly highlight the need to both select patients based on Onc-p53 status and employ rational combinations to truly test the clinical utility of PIs in NSCLC.

*TP53* is often missense mutated within its DNA binding domain (Onc-p53), losing its tumor suppressor role along with acquiring oncogenic functions (also termed Gain of Function in the literature), facilitating tumor cell growth, proliferation, and apoptotic evasion (36). Published studies clearly demonstrated the connection between Onc-p53 and increased proteosome activity in several cancers, including breast, pancreatic, ovarian and prostate (9). Furthermore, NRF2 was shown to complex with Onc-p53 and bind to the promoter region of 26S proteasome subunit genes leading to upregulation of proteosome subunit transcription (9,37). Since Onc-p53 has a high tendency to aggregate in cells (38,39), it has been hypothesized that proteotoxic stress from Onc-p53 aggregates may induce proteasome activity to maintain cellular homeostasis and manage the proteotoxic load (40). Indeed, the p53 R273H allele expressed in H1975 cells has been observed to form intracellular aggregates in breast and ovarian cancer cells using an antibody sensitive to high molecular weight protein aggregates, though the aggregation status of p53 R267P (H1437 cells) has not yet been reported (39,41). Though we did not explore 26S proteasome subunit transcription in this paper, our data indicates that Onc-p53 causes basal proteotoxic stress as indicated by elevated basal proteasome activities in Onc-p53 NSCLC cells, and this stress can be linked through basal ROS generation (13) to activation of NRF2-dependent transcriptional programs, including NRF2-dependent proteasome subunit transactivation, suggesting a positive feedback loop may sustain proteasome gene expression in Onc-p53 NSCLC cells (9). Unlike prior reports in other solid tumor settings, such as breast cancer (9), our work for the first-time links proteotoxic stress generated by Onc-p53 to heightened PI sensitivity, which to date has only been noted in B-cell malignancies such as multiple myeloma and mantle cell lymphoma, where immunoglobulin synthesis causes substantial and ongoing proteotoxic stress (11).

In agreement with published studies, Onc-p53 NSCLC cells demonstrate higher basal levels of ROS (and lower glutathione levels) compared to WT p53 NSCLC cells (**Fig. 2**). Our work, for the first time, also links additional ROS induced by PIs to a cytotoxic cascade that includes ROS-dependent NRF2 activation, downstream activation of ATF3 then NOXA, followed by apoptosis (see **Fig. 6**). Indeed, every step in this cascade can be abrogated by inclusion of the antioxidant NAC or GSH-mimetic GSH-EE during exposure to PI (**Figs. 2F, 3H, 4A-C, S1A, S2C-G**). What remains unclear is the exact source of both basal and PI-induced ROS. One possible explanation is glutathione depletion through means other than consumption by ROS, such as downregulation of transporter or biosynthetic genes regulating the glutathione synthesis pathway (42). Another source of ROS could be oxidative stress due to Onc-p53 itself, either via an induced downstream oxidative stress pathway or through oxidative activity of proteasomes, whose activity is upregulated by Onc-p53 (**Figs. 1A-B and 6**). Further investigation of the underlying mechanism of toxic ROS generation by PIs in Onc-p53 NSCLC could lead to identification of novel targets and improved combination therapies that potentiate the selective cancer cell cytotoxicity of PIs in Onc-p53 NSCLC cells and tumors.

**Fig. 6.**
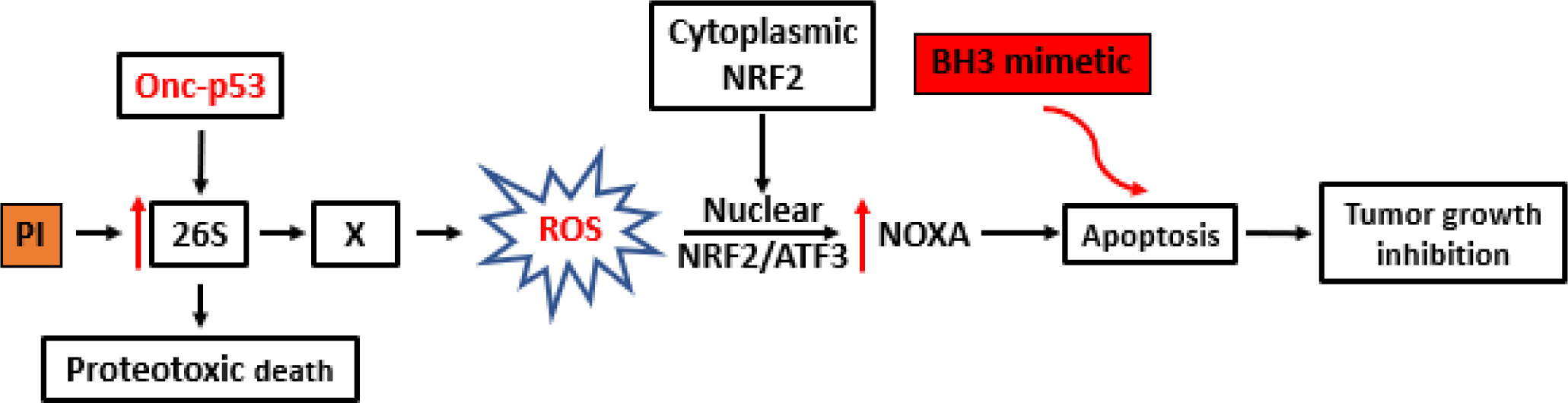
Proposed cytotoxic mechanism of proteasomal inhibitors (PIs) and sensitizing agents in Onc-p53 NSCLC cells. Onc-p53 causes disturbed oxidative homeostasis in NSCLC cells through upregulation of the 26S proteasome and possibly accumulation of misfolded aggregated Onc-p53 protein. PIs further disrupt oxidative homeostasis via an unknown factor or process (“X”) in Onc-p53 NSCLC cells leading to induction and nuclear translocation of NRF2 (after release from the cytoplasmic KEAP1 complex), followed by NRF2-dependent induction of ATF3, which drives expression of the BH3-only protein NOXA, which induces apoptotic cell death that can be sensitized by BH3-mimetics.

NRF2 is a well-established regulator of antioxidant mechanisms whose activity is normally suppressed due to ubiquitination and proteasomal degradation caused by cytoplasmic interaction with the E3 ligase KEAP1 (43). Upon cellular exposure to oxidative stress, NRF2 is released from KEAP1 and translocates to the nucleus to activate highly context and cell-type specific transcriptional programs that broadly regulate cellular defenses to oxidation but also impact both oncogenic and tumor suppressive pathways associated with the hallmarks of cancer (43). Of note, the vast majority of Onc-p53 tumors and cells maintain wild-type KEAP1 (17), though NRF2 can still be induced in the presence of excess oxidative stress that releases it from KEAP1, both stabilizing NRF2 and allowing it to translocate to the nucleus and activate downstream target genes (44). Notably, we observed accumulation and nuclear relocalization of NRF2 protein in Onc-p53 cells upon PI treatment, and both of these effects were blocked by NAC or p53 knockout (**Figs. 4A-C, 4F**). Given prior reports in other settings that oxidative stress causes dissociation of the KEAP1/NRF2 complex, we hypothesize that NRF2 accumulation and nuclear relocalization are due to ROS-induced KEAP1/NRF2 dissociation and NRF2 stabilization, as opposed to direct stabilization of NRF2 by PI treatment or induction of NRF2 transcription (44).

Beyond oxidative activation of NRF2, others have also reported that Onc-p53 can form a complex with and coactivate NRF2 as an alternative or additional means of NRF2 activation in Onc-p53 cancer cells (9,37). Given the obvious complexity of NRF2 regulation both in normal cells and Onc-p53 cancer cells, a more detailed understanding of the mechanism of NRF2 regulation after PI treatment of NSCLC cells will require further study beyond the scope of this work.

The induction of NOXA by PIs has been well established in multiple myeloma, mantle cell lymphoma, chronic myelogenous leukemia, and melanoma (15,45,46), though the exact mechanism is not entirely clear. In some studies, PI’s induced NOXA protein accumulation via protein stabilization through inhibition of the ubiquitin-proteasome pathway (47). In other contexts, however, *NOXA* induction by PIs occurred at the transcriptional level (46). Mechanistically, c-MYC has been reported to mediate PI induction of *NOXA* in melanoma cells (48), while *NOXA* transcription can also be regulated by ATF3/ATF4 in the non-PI setting through a variety of signaling pathways, including the ER stress pathway (49). In Onc-p53 NSCLC cells, we observed BTZ-mediated ATF3, but not ATF4 induction (**Figs. 3G-H, and S2A-B**), which heretofore has not been observed after PI treatment in other tumor contexts, such as multiple myeloma (50). Furthermore, downregulation of ATF3 significantly inhibited BTZ-induction of NOXA and cell death (**Figs. 3I-J**). We and others have shown that ATF3 is capable of binding to the NOXA promoter in other tumor contexts (e.g. squamous cell carcinoma of the head and neck) after other exposures, such as cisplatin (51,52). In the current context with BTZ treatment in Onc-p53 NSCLC cells, we speculate that BTZ-induced ATF3 forms homodimers and binds to the NOXA promoter directly to activate its expression, which we are currently exploring.

Given that BTZ is potently cytotoxic in cultured Onc-p53 NSCLC cells but appears of limited efficacy as a single agent both in our xenograft mouse model (**Figs. 5C and 5H**) and in unselected patients in clinical trials (10), we hypothesized that a combination of BTZ with an agent that abrogates pro-survival signaling, such as a BH3-mimetic (53,54), would enhance BTZ efficacy. Specifically, NOXA’s proapoptotic function is to inactivate pro-survival MCL-1, and while BCL-X_L_ is robustly expressed in both H1975 and H1437 cells, BCL-2 is expressed in H1975 cells but is undetectable in BTZ-sensitive H1437 cells (**Fig. S4A**), suggesting that a BH3-mimetic that can target BCL-X_L_ such as the dual BCL-2/BCL-X_L_ inhibitor navitoclax, may further enhance cell death in Onc-p53 H1975 and H1437 cells already sensitized to BTZ cytotoxicity. Indeed, we observed significant and possibly synergistic enhancement of cell death after BTZ/navitoclax combination treatment in both cell lines *in vitro* (**Figs. 5E and S4B**). We further demonstrated significant tumor growth attenuation by BTZ/navitoclax combination treatment in the H1975 tumor model (**Fig. 5H**). However, navitoclax demonstrates on-target severe inhibition of BCL-X_L_ in platelets, inducing clinically unacceptable thrombocytopenia (27,28), though this was not observed in our mouse xenograft experiment.

Recognizing the hematologic toxicity of targeting BCL-X_L_, next generation BH3-mimetic drugs in development include DT-2216, a PROTAC that is specifically activated to degrade BCL-X_L_ in tumor cells but not in platelets, and mirzotamab clezutoclax (ABBV-155), an antibody-drug conjugate BCL-X_L_ inhibitor (55,56). Both drugs are currently undergoing evaluation in clinical trials (e.g., NCT04886622, NCT03595059) and either could replace navitoclax and serve as a suitable partner for future combination clinical trials with PIs in Onc-p53 NSCLC.

Taken together, this work demonstrates a novel mechanism of action for PIs in Onc-p53 NSCLC cells that has not been observed in tumor types where PIs are currently in clinical use. Our work points the way to potential clinical repurposing of PIs in Onc-p53 NSCLC, and possibly other high frequency Onc-p53 tumors (small cell lung cancer, ovarian cancer, pancreatic cancer, among others). We show enhanced effectiveness of BTZ when combined with carboplatin *in vitro* or in an *in vivo* xenograft model, suggesting that combination of BTZ with standard of care chemotherapy may show effectiveness in selected Onc-p53 NSCLC patients. We did not determine if carboplatin enhanced any specific aspect of the BTZ mechanism of action, though it is known that platinum agents generally increase oxidative stress, and thus may drive enhanced NRF2 activation upon BTZ exposure (57).

By carefully mapping the pathway by which PIs kill Onc-p53 NSCLC cells via proteotoxic stress inducing oxidative stress, followed by induction of an NRF2-ATF3-NOXA signaling cascade and ultimately, apoptosis, we have also developed a rational combination of BTZ with navitoclax as a proof of principle for a future clinical trial involving a PI/BH3-mimetic combination, possibly employing the exciting and lower toxicity BCL-X_L_ PROTAC, DT-2216. Additional translational consideration can be given to PI combinations with agents that target proteotoxic stress (e.g. p97 inhibitors (58)) or agents other than chemotherapy that augment oxidative stress and glutathione depletion (59), such as sulfasalazine and APR-246/PRIMA-1 (9,42,60,61).

## Acknowledgements

The authors thank D. Gewirtz for providing H460 and H460-p53KO cell lines and N. Luffman for technical assistance. Services and products in support of the research project were generated by the Virginia Commonwealth University Cancer Mouse Models Core Laboratory, supported, in part, with funding to VCU Massey Cancer Center from NIH-NCI Cancer Center Support Grant P30 CA016059 and by the USC Norris Flow Cytometry Core supported, in part, by NIH-NCI Cancer Center Support Grant P30 CA014089. This work was supported by VCU Massey Comprehensive Cancer Center Team Science Award.

## Materials and Methods

### Cell lines and Drugs

A549 cells were purchased from ATCC (Manassas, VA, USA). H1975 and H1437 cell lines were the generous gift of S. Deb (VCU), and H460 and H460-p53KO cells (16) were the generous gift of D. Gewirtz (VCU) and all were verified by short tandem repeat (STR) polymorphism analysis. All cell lines were maintained in RPMI 1640 media (Thermo Fisher, 11875093, Waltham, MA, USA), supplemented with 10% heat-inactivated fetal bovine serum (FBS) (Corning, 35-010-CV, Corning, NY, USA) and 100 µg/ml penicillin G/streptomycin at 37°C in a humidified, 5% CO_2_ incubator. Bortezomib (MedChemExpress HY-10227, Monmouth, NJ, USA, and LC Laboratories B-1408, Woburn, MA, USA), carfilzomib (Selleck Chemicals, S2853, Houston, TX, USA), navitoclax (MedChemExpress HY-10087), QVD-O-Ph (MedChemExpress, HY-12305), N-acetyl-l-cysteine (NAC) (MedChemExpress, HY-134495), and glutathione ethyl ester (GSH-EE) (MedChemExpress, HY-134124) were dissolved in dimethyl sulfoxide (DMSO); carboplatin (MedChemExpress, HY-17393, Selleck Chemicals, S1215, Houston, TX, USA) was dissolved in H_2_O and stable drugs were stored at -20°C in the dark. The final concentration of DMSO was 0.1%.

### Generation of cell lines expressing shRNA, cDNA and CRISPR/Cas9 knockouts

Lentiviral short-hairpin RNA (shRNA) vectors, shNOXA (TCRN0000338867), shNOXA#5 TCRN0000338864), shNRF2 (TCRN0000273494), shATF3 (TCRN0000329690), and shATF3#3 (TCRN000013568) were purchased from Sigma Aldrich (St. Louis, MO, USA). shControl (1864), GFP (35637) and p53R273H expression vectors (22934) were purchased from Addgene (Watertown, MA, USA). Each plasmid was co-transfected into HEK293T cells with psPAX2 (Addgene, 12260) and pMD2.G (Addgene, 12259) using Endofectin (GeneCopoeia, EF001, Rockville, MD, USA). Lentivirus-containing supernatants were collected and were used to infect the cell line of interest, and stable cell lines were established by either 2 µg/mL puromycin or 1.0 mg/mL G418 sulfate selection. Control (#2) and p53 gRNA (#4) vectors targeting human *TP53* (GenScript, pLentiCRISPR v2, Piscataway, NJ, USA) were used to homozygously delete the *TP53* gene in H1975 cells as per manufacturer instructions.

### Cell Viability Assays

Cells were seeded at a density of 5 x 10^3^ cells/well in 96-well plates with 100 µL of RPMI media. Cells were treated with the indicated concentrations of either bortezomib or carfilzomib alone for 72 h, or in combination with carboplatin or navitoclax for 96 h. Cell viability was analyzed by WST-1 (Sigma, 11644807001) at 450 nm, or using 3-(4,5-dimethylthiazol-2-yl)-2,5-diphenyltetrazolium bromide (MTT) reagent (Fisher Scientific, AC158992500, Waltham, MA, USA) at 570 nm. WST-1 analyses were performed using a microplate reader from Promega, and CLARIOstar Plate Reader for MTT analyses, according to the manufacturer’s protocol.

### 20S proteasome activity

Cells were seeded at a density of 2.5 x 10^4^ cells/well in a 96-well plate. Following the adherence of cells, proteosome loading solution was prepared according to a fluorometric kit (Sigma Aldrich, MAK172) and added to each well. After 1 h of incubation at 37°C in the dark, mean fluorescence intensity was measured using CLARIOstar plate reader at λ_ex_ = 490 nm and λ_em_ = 525 nm.

### Glutathione assay

1.0 x 10^6^ cells were harvested, pelleted by centrifugation, lysed and deproteinated as per the manufacturer specifications. Total glutathione (GSH) levels in the cells were estimated using a glutathione assay kit (Cayman Chemical, 703002, Ann Arbor, MI, USA) by analyzing the absorbance measured after the addition of the assay cocktail reagent mixture to the deproteinated lysate in the 96-well plate. Absorbance was measured at 412 nm after 25 min of incubation in the dark at room temperature using CLARIOstar plate reader. Total GSH concentration in the cells was calculated using the standard curve determined as per the manufacturer instructions.

### ROS assays

To determine absolute ROS levels, cells were seeded at a density of 2.5 x 10^4^ cells/well in 96-well plates. Following the adherence of cells, ROS indicator dye was prepared according to the manufacturer’s protocol using DCFDA/H2DCFDA - Cellular ROS Assay Kit (Abcam, ab113851, Cambridge, UK), and was added to each well. After 30 min of incubation, the fluorescence intensity was determined using spectrophotometer at Ex/Em = 485/535 nm using CLARIOstar plate reader.

To determine relative ROS levels between cell lines, approximately 1.0 x 10^6^ cells were harvested following the treatments and incubated with the ROS indicator dye, CM-H2DCFDA (ThermoFisher Scientific, C6827, Waltham, MA, USA), for 30 min to 1 h at room temperature according to the manufacturer’s instructions. H2DCFDA-stained single cell suspension was run through BD-FACSCanto II flow cytometer (USC Norris Core) at Ex/Em = 492–495/517–527 nm. Percentages of ROS positive single cells (detected by FITC-channel) were recorded using BD FACSDiva 8.0 software.

### Annexin-V-FITC

Cells were seeded at a density of 5.0 x 10^5^ cells in 60mm dishes and were treated with either vehicle (DMSO), 5 nM bortezomib, 1 µM QVD-O-Ph, or in combination for 48 h. Following treatment, cells were harvested, washed with PBS and resuspended with 100 µL of 1x Annexin V binding buffer (BD Biosciences, 556454, Franklin Lakes, NJ, USA). Annexin-V FITC (BioLegend, 640945, San Diego, CA, USA) and propidium iodide (PI) (Thermo Fisher, P3566) were added, and cells were incubated in the dark for 15 min at room temperature. Then, 400 µL of 1x binding buffer were added to the suspension for analysis. Cells were analyzed using FACSCAN (BD FACSCanto) at the VCU Flow Cytometry Core and quantified the population as either double-negative, Annexin V-positive, PI-positive, or double-positive by the FlowJo software version 10.8.1.

### Immunoblot analyses

Cells were seeded at a density of 1.0 x 10^6^ in 10cm dishes. Cells were treated with either vehicle (DMSO), 5 nM bortezomib, 1 µM QVD-O-Ph, 1 mM NAC, 1 mM GSH-EE, or in combination for 48 h. Whole cell lysates were prepared using CHAPS buffer [20 mM Tris (pH 7.4), 137 mM NaCl, 1 mM dithiothreitol (DTT), 1% CHAPS (3-[(3-cholamidopropyl) dimethylammonio] 1-propanesulfonate)]. Equal amounts of protein were loaded into an SDS-polyacrylamide gel, transferred onto a nitrocellulose membrane, incubated with antibodies of interest, and analyzed with ECL2 western blotting substrate (Thermo Scientific, 32132). Primary antibodies were used in a 1:1000 dilution for p53 (Santa Cruz, DO-1, Santa Cruz, CA, USA), GAPDH (Cell Signaling, D64E10, Danvers, MA, USA), Cleaved Caspase-3 (Cell Signaling, D175), NOXA (Invitrogen, MA1-41000, Waltham, MA, USA), ATF3 (Abcam, ab207434), NRF2 (Cell Signaling, D1Z9C). Secondary antibodies were used in a 1:2000 dilution for HRP-linked anti-rabbit IgG (Cell Signaling) and HRP-linked anti-mouse IgG (Cell Signaling).

### Quantitative Reverse Transcriptase Polymerase Chain Reaction (qRT-PCR)

Cells were seeded at a density of 1.0 x 10^6^ in 10cm dishes and were treated with either vehicle (DMSO), 5 nM bortezomib, 1 mM NAC, 1mM GSH-EE, or in combination for 48 h. Cells were harvested and total RNA was extracted using Quick-RNA MiniPrep (Zymo, R1054, Irvine, CA, USA) following the manufacturer’s instructions. cDNA was synthesized using Applied Biosystems High Capacity cDNA reverse transcription kit (Thermo Fisher, 4368814) based on the manufacturer’s protocol. cDNA was amplified in triplicate using PowerTrack SYBR green master mix (Thermo Fisher, A46109) in the StepOnePlus Real-time PCR system by Applied Biosystems. qRT-PCR primers were synthesized with the sequences listed in **Fig S5** and mRNA expression was determined using ΔΔCt.

### Immunofluorescence

Cells were seeded at a density of 5.0 x 10^2^ cells/well in 4-well culture slides and treated with 10 nM bortezomib, 1 mM NAC, or combination for 24 h. Following treatment, cells were fixed by incubating with 4% paraformaldehyde for 30 min at room temperature. After a 2 min wash with 1x PBS, cells were permeabilized and blocked using 0.2% Triton-X in 5% BSA for 30 min at room temperature. Following a wash with 1x PBS, cells were incubated with primary antibody diluted in blocking buffer for 1 h at room temperature. After washing 3-times with 1x PBS, cells were incubated with secondary antibody for 1 h at room temperature. After washing 3-times with 1x PBS, slides were cover slipped using DAPI with mounting media. Fluorescent cells were imaged and analyzed using Leica fluorescent microscope at 20X magnification. Primary antibodies were used in a 1:100 dilution for NRF2 (Santa Cruz, sc-13032). Secondary antibodies were used in a 1:300 dilution for mouse anti-rabbit IgG-CFL 594 (Santa Cruz, sc-516250).

### In Vivo Studies

All animal studies were conducted in accordance with VCU or USC Institutional Animal Care and Use Committee (IACUC) guidelines. For experiments in **Figs. 5C-D**, A549 (5.0 x 10^6^ cells) or H1975 (1.0 x 10^7^ cells) were subcutaneously injected into the flank of male 2-4-month-old NOD-SCID-II.2gamma receptor null (NSG) mice obtained from either the VCU Cancer Mouse Models Core (**Fig. 5D**) or from Jackson Laboratories (**Fig. 5C**). Once tumors became palpable (∼150 mm^3^), mice were randomized into 4 groups (4 mice/group) and were treated with vehicle, BTZ (0.6 mg/kg) 2x/week, carboplatin (25 mg/kg) weekly, and BTZ + carboplatin combination via intraperitoneal injection (IP) for 4 weeks. For the experiment in **Fig. 5H**, H1975 cells (4.0 x 10^6^) were subcutaneously inoculated into the flank of 6-week-old female NSG mice obtained from the VCU Cancer Mouse Models Core. Once tumors became palpable (∼100 mm^3^), mice were randomized into 4 groups (5 mice/group) and were treated with vehicle, navitoclax (80 mg/kg) 3x/week via oral gavage, BTZ (1 mg/kg) 2x/week via IP injection, and BTZ + navitoclax combination for 4 weeks. Tumor volume was calculated as V= ½ x AB^2^, where A is the longest dimension of the tumor and B is the dimension of the tumor perpendicular to A as measured by caliper.

### Immunohistochemistry

Tumors were fixed in 10% formalin phosphate buffer and paraffin embedded. Tissue embedding and sectioning was performed by VCU Cancer Mouse Models Core. Slides were blocked for 20 min in PowerVision Universal IHC Blocking diluent (Leica, PV6123), then stained with cleaved caspase-3 antibodies (Cell Signaling, 9664), at a 1:500 dilution for 45 min. Staining was performed using Leica Bond RX autostainer using Polymer detection system. Images were taken on the Vectra Polaris Automated Quantitative Pathology Imaging System (Akoya Biosciences) at 20X magnification. Scale bar indicates 50 µM.

### Circulating neutrophil/platelet analysis

Immediately after the last day of treatment of the xenograft experiment in **Fig. 5H**, blood samples (∼0.2 ml) were collected by cheek bleeding using EDTA coated syringes and immediately analyzed for neutrophil and platelet counts using a Hemavet 950FS (Drew Scientific, Miami Lakes, FL, USA) in the VCU Cancer Mouse Models Core.

### Statistical analyses

All quantitative data are shown as ± 1.0 S.D. from at least three independent experiments each of which included technical duplicates. All pairwise statistical comparison t-tests were performed using GraphPad Prism Software Version 8.1 and/or Microsoft Excel. p≤0.05 was considered statistically significant.

## Supplementary Figure legends

**Fig. S1. BTZ-induced cell death depends on GSH and NOXA. A.** H1975 cells were treated with vehicle or 5 nM BTZ with or without 1 mM GSH-EE for 48 h. Cell viability was determined by Trypan-blue exclusion assay. \**p*<0.05. Error bars indicate +/- 1.0 S.D. **B.** H1437 cells were treated with the indicated concentration of BTZ for 48 h. Whole cell lysates were immunoblotted with the indicated antibodies. **C.** H1975 cells stably expressing control shRNA (shC) or an alternate NOXA shRNA, shNOXA#5, were treated with vehicle (-) or BTZ (5 nM) for 48 h and cell lysates were immunoblotted with the indicated antibodies. **D.** H1437 shRNA control and shNOXA cells were treated with and without BTZ (5 nM) for 48 h and cell lysates were immunoblotted with the indicated antibodies.

**Fig. S2. Oxidative stress-dependent induction of ATF3 upon BTZ exposure in Onc-p53 NSCLC cells drives NOXA induction and cell death. A.** H1975 cells were treated with vehicle or BTZ (2, 5 nM) for 48 h and mRNA expression of ATF4 was analyzed by qRT-PCR. **B.** H1437 cells were treated with vehicle, 5, or 10 nM of BTZ for 48 h and cell lysates were immunoblotted with ATF3 and GAPDH antibodies. **C-D.** H1975 cells were treated with vehicle or BTZ (5 nM) with or without NAC (1 mM) or for 48 h and NOXA and ATF3 expression were analyzed by qRT-PCR. **E.** H1975 cells were treated with vehicle or BTZ (5 nM) with or without GSH-EE (1mM; GSH) for 48 h and cell lysates were immunoblotted with indicated antibodies. **F-G.** H1975 cells were treated with vehicle or BTZ (5 nM) with or without GSH-EE (1 mM; GSH) for 48 h and NOXA and ATF3 expression were analyzed by qRT-PCR. **H.** H1975 cells stably expressing control shRNA (shC) or an alternate ATF3 shRNA, shATF3#3, were treated with vehicle (-) or BTZ (5 nM) for 48 h and cell lysates were subjected to immunoblotting with the indicated antibodies. \**p*<0.05, \*\**p*<0.01, \*\*\**p*<0.005. Error bars indicate +/- 1.0 S.D.

**Fig. S3. BTZ-mediated ATF3/NOXA induction requires Onc-p53. A-B.** H1975-Control or H1975-p53KO cells were treated with BTZ (5 nM) for 48 h and NOXA and ATF3 expression were analyzed by qRT-PCR. \*\**p*<0.01, \*\*\**p*<0.005. Error bars indicate +/- 1.0 S.D.

**Fig. S4. Navitoclax enhances BTZ-induced cytotoxicity in Onc-p53-expressing NSCLC cells. A.** Basal levels of anti-apoptotic MCL-1, BCL-X_L_ and BCL-2 proteins were determined by immunoblotting. **B**. H1437 cells were treated with vehicle or the indicated concentrations of navitoclax with/without BTZ (5 nM) for 96 h. Cell viability was determined by WST-1 assay. \*\**p*<0.01. Error bars indicate +/- 1.0 S.D. **C**. H1975 tumors treated with BTZ and/or navitoclax in **Fig. 5H** were harvested at the endpoints and stained by IHC with cleaved caspase-3 antibody. The images are representative of 3 independent tumors from each treatment. **D.** The average percentage of cleaved caspase-3 positive cells in **C** were quantified from 5 random fields of each tumor section at 40X magnification. **E.** Circulating platelet and neutrophil counts at the experimental endpoint from mice treated with vehicle, BTZ, navitoclax, or the combination. Normal neutrophil and platelet count ranges are indicated.

**Fig. S5. Primer sequences used for qRT-PCR.**

## References

1. Siegel RL, Giaquinto AN, Jemal A. Cancer statistics, 2024. CA: A Cancer Journal for Clinicians. 2024;74:12–49.

2. Ganti AK, Klein AB, Cotarla I, Seal B, Chou E. Update of Incidence, Prevalence, Survival, and Initial Treatment in Patients With Non–Small Cell Lung Cancer in the US. JAMA Oncology. 2021;7:1824–32.

3. 3. Chen X, Zhang T, Su W, Dou Z, Zhao D, Jin X, et al. Mutant p53 in cancer: from molecular mechanism to therapeutic modulation. Cell Death Dis. Nature Publishing Group; 2022;13:1–14.

4. Mogi A, Kuwano H. TP53 mutations in nonsmall cell lung cancer. J Biomed Biotechnol. 2011;2011:583929.

5. Feng H, Xu H, Shi X, Ding G, Yan C, Li L, et al. TP53 Exon 5 Mutation Indicates Poor Progression-Free Survival for Patients with Stage IV NSCLC. Front Biosci (Landmark Ed). 2023;28:147.

6. Singh S, Vaughan CA, Frum RA, Grossman SR, Deb S, Palit Deb S. Mutant p53 establishes targetable tumor dependency by promoting unscheduled replication. J Clin Invest. 2017;127:1839–55.

7. Skoulidis F, Heymach JV. Co-occurring genomic alterations in non-small-cell lung cancer biology and therapy. Nat Rev Cancer. 2019;19:495–509.

8. Wadowska K, Bil-Lula I, Trembecki Ł, Śliwińska-Mossoń M. Genetic Markers in Lung Cancer Diagnosis: A Review. Int J Mol Sci. 2020;21:E4569.

9. Walerych D, Lisek K, Sommaggio R, Piazza S, Ciani Y, Dalla E, et al. Proteasome machinery is instrumental in a common gain-of-function program of the p53 missense mutants in cancer. Nat Cell Biol. 2016;18:897–909.

10. Oduah EI, Grossman SR. Harnessing the vulnerabilities of p53 mutants in lung cancer – Focusing on the proteasome: a new trick for an old foe? Cancer Biol Ther. 21:293–302.

11. Ho Zhi Guang M, Kavanagh EL, Dunne LP, Dowling P, Zhang L, Lindsay S, et al. Targeting Proteotoxic Stress in Cancer: A Review of the Role that Protein Quality Control Pathways Play in Oncogenesis. Cancers (Basel). 2019;11:66.

12. Manasanch EE, Orlowski RZ. Proteasome Inhibitors in Cancer Therapy. Nat Rev Clin Oncol. 2017;14:417– 33.

13. Cordani M, Butera G, Pacchiana R, Masetto F, Mullappilly N, Riganti C, et al. Mutant p53-Associated Molecular Mechanisms of ROS Regulation in Cancer Cells. Biomolecules. 2020;10:361.

14. Aggarwal V, Tuli HS, Varol A, Thakral F, Yerer MB, Sak K, et al. Role of Reactive Oxygen Species in Cancer Progression: Molecular Mechanisms and Recent Advancements. Biomolecules. 2019;9:735.

15. Pérez-Galán P, Roué G, Villamor N, Montserrat E, Campo E, Colomer D. The proteasome inhibitor bortezomib induces apoptosis in mantle-cell lymphoma through generation of ROS and Noxa activation independent of p53 status. Blood. 2006;107:257–64.

16. Patel NH, Xu J, Saleh T, Wu Y, Lima S, Gewirtz DA. Influence of nonprotective autophagy and the autophagic switch on sensitivity to cisplatin in non-small cell lung cancer cells. Biochem Pharmacol. 2020;175:113896.

17. DepMap: The Cancer Dependency Map Project at Broad Institute [Internet]. [cited 2023 Dec 30]. Available from: https://depmap.org/portal/

18. Gomez-Bougie P, Wuilleme-Toumi S, Menoret E, Trichet V, Robillard N, Philippe M, et al. Noxa Up-regulation and Mcl-1 Cleavage Are Associated to Apoptosis Induction by Bortezomib in Multiple Myeloma. Cancer Res. 2007;

19. Pietkiewicz S, Sohn D, Piekorz RP, Grether-Beck S, Budach W, Sabapathy K, et al. Oppositional Regulation of Noxa by JNK1 and JNK2 during Apoptosis Induced by Proteasomal Inhibitors. PLOS ONE. Public Library of Science; 2013;8:e61438.

20. Bi Z, Fu Y, Wadgaonkar P, Qiu Y, Almutairy B, Zhang W, et al. New Discoveries and Ambiguities of Nrf2 and ATF3 Signaling in Environmental Arsenic-Induced Carcinogenesis. Antioxidants (Basel). 2021;11:77.

21. Kha M-L, Hesse L, Deisinger F, Sipos B, Röcken C, Arlt A, et al. The antioxidant transcription factor Nrf2 modulates the stress response and phenotype of malignant as well as premalignant pancreatic ductal epithelial cells by inducing expression of the ATF3 splicing variant ΔZip2. Oncogene. Nature Publishing Group; 2019;38:1461–76.

22. Kim K-H, Jeong J-Y, Surh Y-J, Kim K-W. Expression of stress-response ATF3 is mediated by Nrf2 in astrocytes. Nucleic Acids Res. 2010;38:48–59.

23. Ngo V, Duennwald ML. Nrf2 and Oxidative Stress: A General Overview of Mechanisms and Implications in Human Disease. Antioxidants (Basel). 2022;11:2345.

24. Ma Q. Role of Nrf2 in Oxidative Stress and Toxicity. Annu Rev Pharmacol Toxicol. 2013;53:401–26.

25. Al-Fahdawi MQ, Al-Doghachi FAJ, Abdullah QK, Hammad RT, Rasedee A, Ibrahim WN, et al. Oxidative stress cytotoxicity induced by platinum-doped magnesia nanoparticles in cancer cells. Biomedicine & Pharmacotherapy. 2021;138:111483.

26. Chiou J-T, Huang N-C, Huang C-H, Wang L-J, Lee Y-C, Shi Y-J, et al. NOXA-mediated degradation of MCL1 and BCL2L1 causes apoptosis of daunorubicin-treated human acute myeloid leukemia cells. J Cell Physiol. 2021;236:7356–75.

27. Schoenwaelder SM, Jarman KE, Gardiner EE, Hua M, Qiao J, White MJ, et al. Bcl-xL-inhibitory BH3 mimetics can induce a transient thrombocytopathy that undermines the hemostatic function of platelets. Blood. 2011;118:1663–74.

28. Kaefer A, Yang J, Noertersheuser P, Mensing S, Humerickhouse R, Awni W, et al. Mechanism-based pharmacokinetic/pharmacodynamic meta-analysis of navitoclax (ABT-263) induced thrombocytopenia. Cancer Chemother Pharmacol. 2014;74:593–602.

29. NOD SCID Mouse Clinical Pathology Data | Charles River (criver.com) [Internet]. [cited 2024 Feb 19]. Available from: https://www.criver.com/sites/default/files/resources/doc_a/NODSCIDMouseClinicalPathologyData.pdf

30. Robak P, Robak T. Bortezomib for the Treatment of Hematologic Malignancies: 15 Years Later. Drugs R D. 2019;19:73–92.

31. Chua ADW, Thaarun T, Yang H, Lee ARYB. Proteasome inhibitors in the treatment of nonsmall cell lung cancer: A systematic review of clinical evidence. Health Science Reports. 2023;6:e1443.

32. Fanucchi MP, Fossella FV, Belt R, Natale R, Fidias P, Carbone DP, et al. Randomized phase II study of bortezomib alone and bortezomib in combination with docetaxel in previously treated advanced non-small-cell lung cancer. J Clin Oncol. 2006;24:5025–33.

33. Davies AM, Chansky K, Lara PN, Gumerlock PH, Crowley J, Albain KS, et al. Bortezomib plus gemcitabine/carboplatin as first-line treatment of advanced non-small cell lung cancer: a phase II Southwest Oncology Group Study (S0339). J Thorac Oncol. 2009;4:87–92.

34. Scagliotti GV, Germonpré P, Bosquée L, Vansteenkiste J, Gervais R, Planchard D, et al. A randomized phase II study of bortezomib and pemetrexed, in combination or alone, in patients with previously treated advanced non-small-cell lung cancer. Lung Cancer. 2010;68:420–6.

35. Davies AM, Ruel C, Lara PN, Lau DH, Gumerlock PH, Bold R, et al. The proteasome inhibitor bortezomib in combination with gemcitabine and carboplatin in advanced non-small cell lung cancer: a California Cancer Consortium Phase I study. J Thorac Oncol. 2008;3:68–74.

36. Alvarado-Ortiz E, de la Cruz-López KG, Becerril-Rico J, Sarabia-Sánchez MA, Ortiz-Sánchez E, García-Carrancá A. Mutant p53 Gain-of-Function: Role in Cancer Development, Progression, and Therapeutic Approaches. Front Cell Dev Biol. 2021;8:607670.

37. Lisek K, Walerych D, Del Sal G. Mutant p53–Nrf2 axis regulates the proteasome machinery in cancer. Mol Cell Oncol. 2016;4:e1217967.

38. Ano Bom APD, Rangel LP, Costa DCF, De Oliveira GAP, Sanches D, Braga CA, et al. Mutant p53 Aggregates into Prion-like Amyloid Oligomers and Fibrils. Journal of Biological Chemistry. 2012;287:28152–62.

39. Levy CB, Stumbo AC, Ano Bom APD, Portari EA, Carneiro Y, Silva JL, et al. Co-localization of mutant p53 and amyloid-like protein aggregates in breast tumors. The International Journal of Biochemistry & Cell Biology. 2011;43:60–4.

40. Jones CL, Tepe JJ. Proteasome Activation to Combat Proteotoxicity. Molecules. 2019;24:2841.

41. Heinzl N, Koziel K, Maritschnegg E, Berger A, Pechriggl E, Fiegl H, et al. A comparison of four technologies for detecting p53 aggregates in ovarian cancer. Front Oncol. 2022;12:976725.

42. Liu DS, Duong CP, Haupt S, Montgomery KG, House CM, Azar WJ, et al. Inhibiting the system xC-/glutathione axis selectively targets cancers with mutant-p53 accumulation. Nat Commun. 2017;8:14844.

43. de la Vega MR, Chapman E, Zhang DD. NRF2 and the hallmarks of cancer. Cancer Cell. 2018;34:21–43.

44. Bellezza I, Giambanco I, Minelli A, Donato R. Nrf2-Keap1 signaling in oxidative and reductive stress. Biochimica et Biophysica Acta (BBA) - Molecular Cell Research. 2018;1865:721–33.

45. Fernandez Y, Verhaegen M, Miller TP, Rush JL, Steiner P, Opipari AW, et al. Differential Regulation of Noxa in Normal Melanocytes and Melanoma Cells by Proteasome Inhibition: Therapeutic Implications. Cancer Res. 2005;

46. Baou M, Kohlhaas SL, Butterworth M, Vogler M, Dinsdale D, Walewska R, et al. Role of NOXA and its ubiquitination in proteasome inhibitor-induced apoptosis in chronic lymphocytic leukemia cells. Haematologica. 2010;95:1510–8.

47. Craxton A, Butterworth M, Harper N, Fairall L, Schwabe J, Ciechanover A, et al. NOXA, a sensor of proteasome integrity, is degraded by 26S proteasomes by an ubiquitin-independent pathway that is blocked by MCL-1. Cell Death Differ. 2012;19:1424–34.

48. Nikiforov MA, Riblett M, Tang W-H, Gratchouck V, Zhuang D, Fernandez Y, et al. Tumor cell-selective regulation of NOXA by c-MYC in response to proteasome inhibition. Proc Natl Acad Sci U S A. 2007;104:19488–93.

49. Wang Q, Mora-Jensen H, Weniger MA, Perez-Galan P, Wolford C, Hai T, et al. ERAD inhibitors integrate ER stress with an epigenetic mechanism to activate BH3-only protein NOXA in cancer cells. Proc Natl Acad Sci U S A. 2009;106:2200–5.

50. Narita T, Ri M, Masaki A, Mori F, Ito A, Kusumoto S, et al. Lower expression of activating transcription factors 3 and 4 correlates with shorter progression-free survival in multiple myeloma patients receiving bortezomib plus dexamethasone therapy. Blood Cancer J. 2015;5:e373.

51. Sharma K, Vu T-T, Cook W, Naseri M, Zhan K, Nakajima W, et al. p53-independent Noxa induction by cisplatin is regulated by ATF3/ATF4 in head and neck squamous cell carcinoma cells. Mol Oncol. 2018;12:788–98.

52. Núñez-Vázquez S, Sánchez-Vera I, Saura-Esteller J, Cosialls AM, Noisier AFM, Albericio F, et al. NOXA upregulation by the prohibitin-binding compound fluorizoline is transcriptionally regulated by integrated stress response-induced ATF3 and ATF4. FEBS J. 2021;288:1271–85.

53. Ni Chonghaile T, Letai A. Mimicking the BH3 domain to kill cancer cells. Oncogene. 2008;27 Suppl 1:S149–157.

54. Diepstraten ST, Anderson MA, Czabotar PE, Lessene G, Strasser A, Kelly GL. The manipulation of apoptosis for cancer therapy using BH3-mimetic drugs. Nat Rev Cancer. 2022;22:45–64.

55. Khan S, Zhang X, Lv D, Zhang Q, He Y, Zhang P, et al. A selective BCL-XL PROTAC degrader achieves safe and potent antitumor activity. Nat Med. 2019;25:1938–47.

56. Tolcher AW, Carneiro BA, Dowlati A, Abdul Razak AR, Chae YK, Villella JA, et al. A first-in-human study of mirzotamab clezutoclax as monotherapy and in combination with taxane therapy in relapsed/refractory solid tumors: Dose escalation results. JCO. Wolters Kluwer; 2021;39:3015–3015.

57. Weniger MA, Rizzatti EG, Perez-Galan P, Liu D, Wang Q, Munson PJ, et al. Treatment-induced oxidative stress and cellular antioxidant capacity determine response to bortezomib in mantle cell lymphoma. Clin Cancer Res. 2011;17:5101–12.

58. Auner HW, Moody AM, Ward TH, Kraus M, Milan E, May P, et al. Combined Inhibition of p97 and the Proteasome Causes Lethal Disruption of the Secretory Apparatus in Multiple Myeloma Cells. PLoS One. 2013;8:e74415.

59. Du Z-X, Zhang H-Y, Meng X, Guan Y, Wang H-Q. Role of oxidative stress and intracellular glutathione in the sensitivity to apoptosis induced by proteasome inhibitor in thyroid cancer cells. BMC Cancer. 2009;9:56.

60. Khoral P, Guo RJ, Abdi J, Chang H. Prima-1Met Combined with Bortezomib Has Synergistic Anti-Myeloma Activity By Modulation of Apoptosis and Cell Cycle Regulating Genes. Blood. 2015;126:4213.

61. Starheim KK, Holien T, Misund K, Johansson I, Baranowska KA, Sponaas A-M, et al. Intracellular glutathione determines bortezomib cytotoxicity in multiple myeloma cells. Blood Cancer Journal. Nature Publishing Group; 2016;6:e446–e446.

