## Supplemental Figures for "Oncogenic Mutant p53 Sensitizes Non-Small Cell Lung Cancer Cells to Proteasome Inhibition via Oxidative Stress-Dependent Induction of Mitochondrial Apoptosis"

### Figure S1

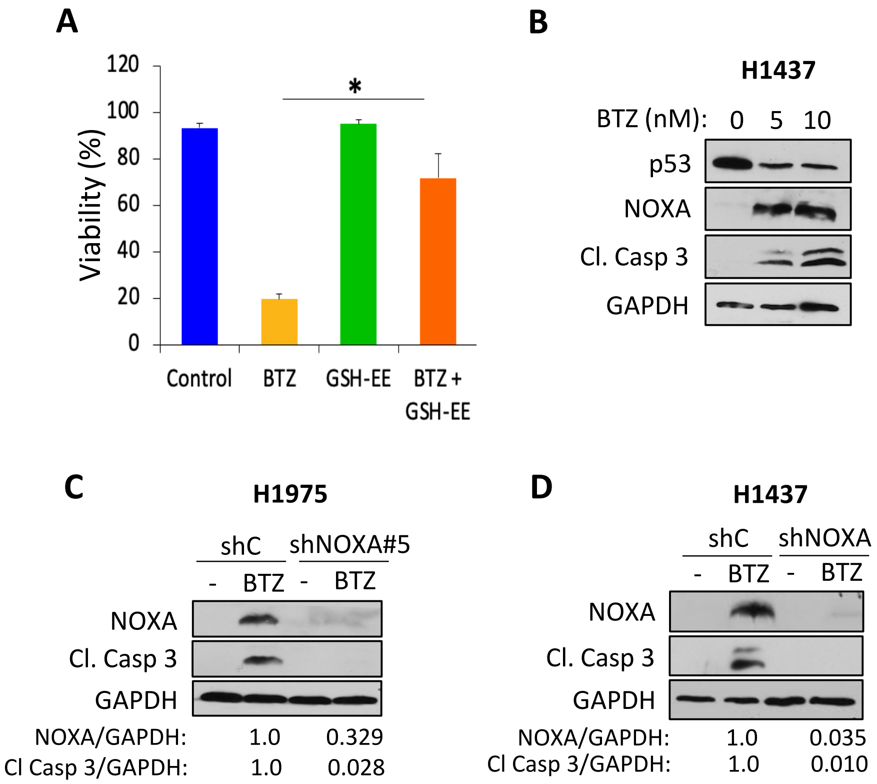

A
120
100
80
60
40
20
0
Viability (%)
*
Control
BTZ
GSH-EE
BTZ + GSH-EE
B
H1437
BTZ (nM): 0 5 10
p53
NOXA
Cl. Casp 3
GAPDH
C
H1975
shC
shNOXA#5
- BTZ - BTZ
NOXA
Cl. Casp 3
GAPDH
NOXA/GAPDH: 1.0 0.329
Cl Casp 3/GAPDH: 1.0 0.028
D
H1437
shC
shNOXA
- BTZ - BTZ
NOXA
Cl. Casp 3
GAPDH
NOXA/GAPDH: 1.0 0.035
Cl Casp 3/GAPDH: 1.0 0.010

**Figure S2**

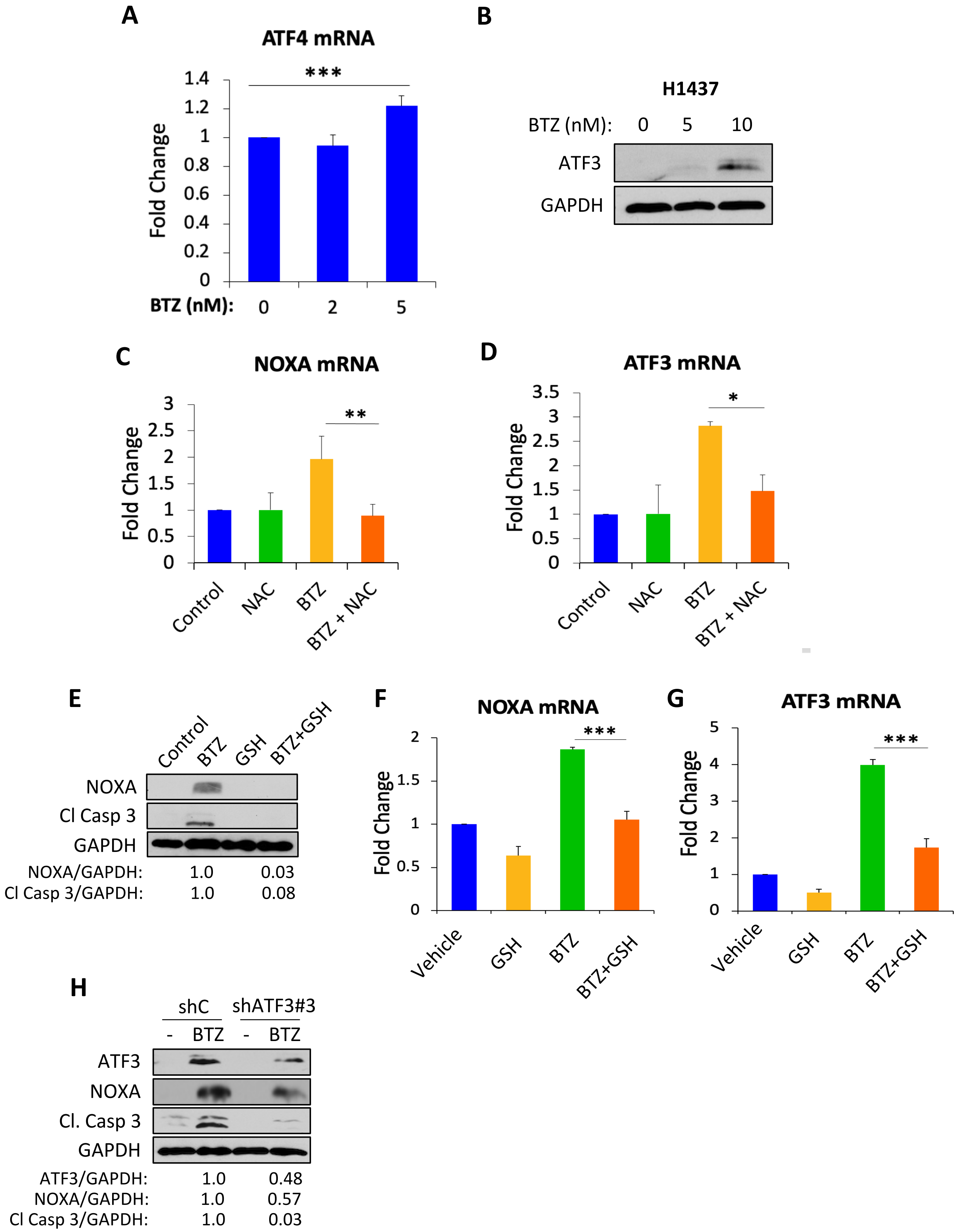
A
ATF4 mRNA
***
Fold Change
1.4
1.2
1
0.8
0.6
0.4
0.2
0
BTZ (nM): 0 2 5
B
H1437
BTZ (nM): 0 5 10
ATF3
GAPDH
C
NOXA mRNA
**
Fold Change
3
2.5
2
1.5
1
0.5
0
Control
NAC
BTZ
BTZ + NAC
D
ATF3 mRNA
*
Fold Change
3.5
3
2.5
2
1.5
1
0.5
0
Control
NAC
BTZ
BTZ + NAC
E
Control
BTZ
GSH
BTZ+GSH
NOXA
Cl Casp 3
GAPDH
NOXA/GAPDH: 1.0 0.03
Cl Casp 3/GAPDH: 1.0 0.08
F
NOXA mRNA
***
Fold Change
2
1.5
1
0.5
0
Vehicle
GSH
BTZ
BTZ+GSH
G
ATF3 mRNA
***
Fold Change
5
4
3
2
1
0
Vehicle
GSH
BTZ
BTZ+GSH
H
shC
shATF3#3
- BTZ - BTZ
ATF3
NOXA
Cl. Casp 3
GAPDH
ATF3/GAPDH: 1.0 0.48
NOXA/GAPDH: 1.0 0.57
Cl Casp 3/GAPDH: 1.0 0.03

**Figure S3**

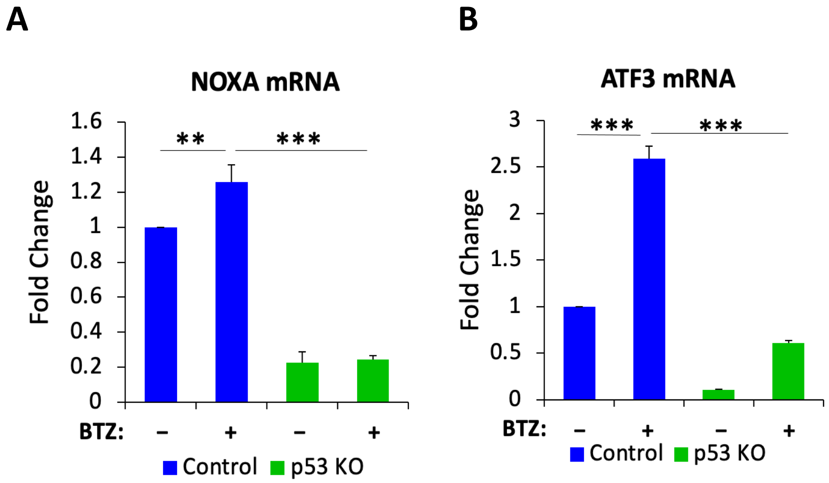
A
NOXA mRNA
Fold Change
1.6
1.4
1.2
1
0.8
0.6
0.4
0.2
0
**
***
BTZ:
–
+
–
+
Control
p53 KO
B
ATF3 mRNA
Fold Change
3
2.5
2
1.5
1
0.5
0
***
***
BTZ:
–
+
–
+
Control
p53 KO

**Figure S4**

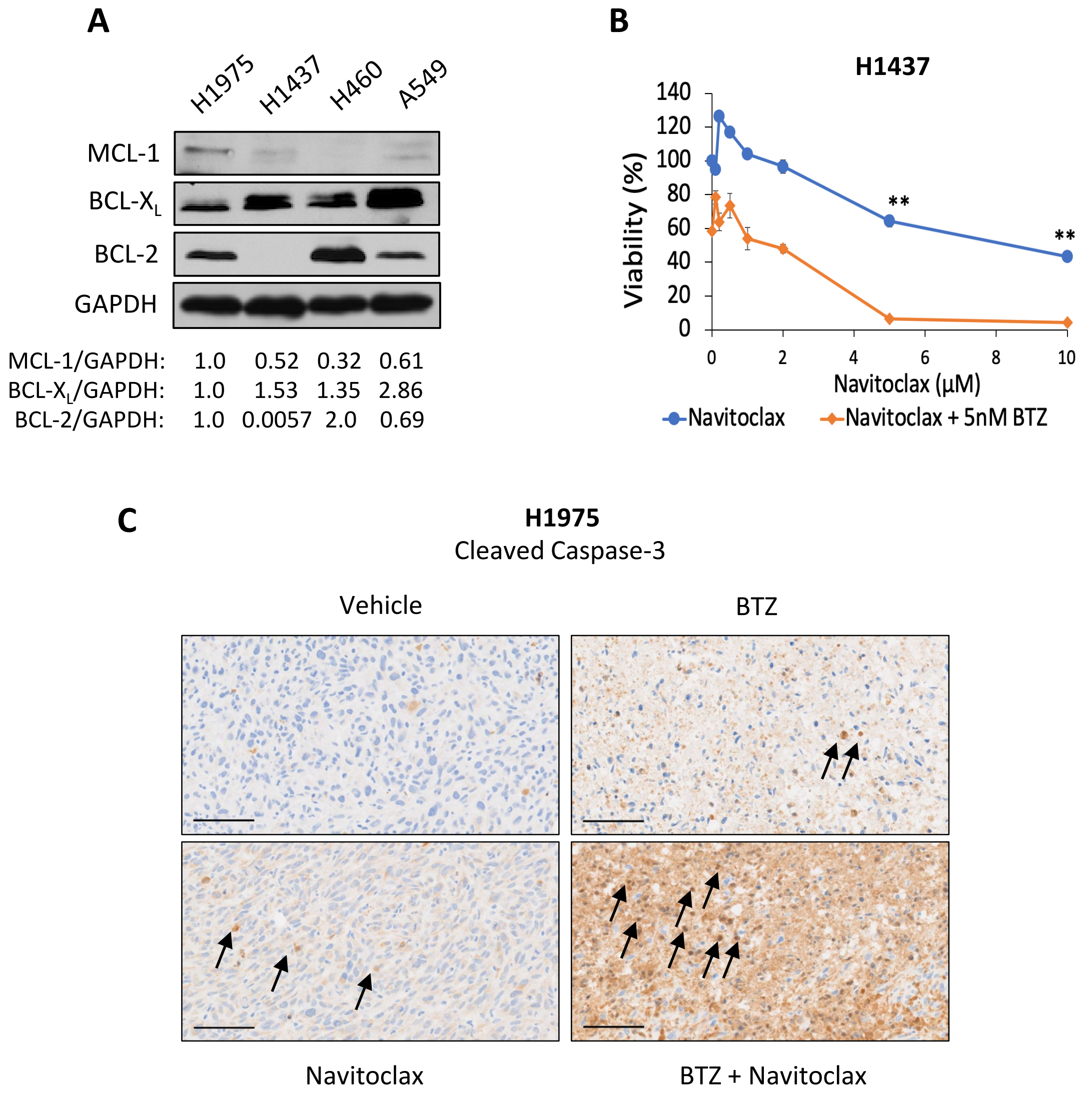


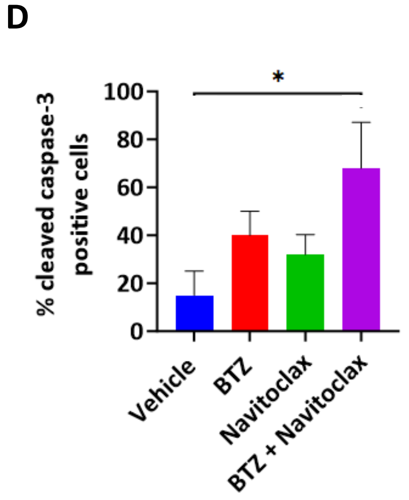


**E**

| | **Neutrophils (K/µL)** (Mean ± SD) | **Platelets (K/µL)** (Mean ± SD) |
|---|---|---|
| **Vehicle** | 1.37 ± 1.12 | 558.67 ± 389.03 |
| **BTZ** | 2.18 ± 1.25 | 797.00 ± 425.79 |
| **Navitoclax** | 1.89 ± 0.42 | 1066.50 ± 232.64 |
| **Navitoclax + BTZ** | 2.76 ± 0.60 | 1347.67 ± 295.74 |
| **Normal range** | 0.6-3.16 | 651-1878 |

### Figure S5

| Gene | Forward Sequence | Reverse Sequence |
| --- | --- | --- |
| ATF3 | 5’- CGC TGG AAT CAG TCA CTG TCA G -3’ | 5’- CTT GTT TCG GCA CTT TGC AGC TG -3’ |
| ATF4 | 5’- TTC TCC AGC GAC AAG GCT AAG G -3’ | 5’- CTC CAA CAT CCA ATC TGT CCC G -3’ |
| NOXA | 5’- CTG GAA GTC GAG TGT GCT ACT C -3’ | 5’- TGA AGG AGT CCC CTC ATG CAA G -3’ |
| p53 | 5’- AAG GAA ATT TGC GTG TGG AGT -3’ | 5’- AAA GCT GTT CCG TCC CAG TA -3’ |
| GAPDH | 5’- GTC TCC TCT GAC TTC AAC AGC G -3’ | 5’- ACC ACC CTG TTG CTG TAG CCA A -3’ |
